# Escaping the Fate of Sisyphus: Assessing Resistome Hybridization Baits for Antimicrobial Resistance Gene Capture

**DOI:** 10.1101/2021.07.20.452950

**Authors:** Megan S. Beaudry, Jesse C. Thomas, Rodrigo Baptista, Amanda Sullivan, William Norfolk, Alison Devault, Jacob Enk, Troy J. Kieran, Olin Rhodes, Allison Perry, Laura Rose, Natalia J. Bayona-Vásquez, Ade Oladeinde, Erin Lipp, Susan Sanchez, Travis C. Glenn

## Abstract

Finding, characterizing, and monitoring reservoirs for antimicrobial resistance (AMR) is vital to protecting public health. Hybridization capture baits are an accurate, sensitive, and cost-effective technique used to enrich and characterize DNA sequences of interest, including antimicrobial resistance genes (ARGs), in complex environmental samples. We demonstrate the continued utility of a set of 19,933 hybridization capture baits designed from the Comprehensive Antibiotic Resistance Database (CARD)v1.1.2 and Pathogenicity Island Database (PAIDB)v2.0, targeting 3,565 unique nucleotide sequences that confer resistance. We demonstrate the efficiency of our bait set on a custom-made resistance mock community and complex environmental samples to increase the proportion of on-target reads as much as >200-fold. However, keeping pace with newly discovered ARGs poses a challenge when studying AMR, because novel ARGs are continually being identified and would not be included in bait sets designed prior to discovery. We provide imperative information on how our bait set performs against CARDv3.3.1, as well as a generalizable approach for deciding when and how to update hybridization capture bait sets. This research encapsulates the full life cycle of baits for hybridization capture of the resistome from design and validation (both *in silico* and *in vitro*) to utilization and forecasting updates and retirement.

**Originality-Significance Statement:** This work is applicable to a wide range of research. It helps to define conditions under which hybridization capture is useful regarding not only antimicrobial resistance specifically, but also more generally how to assess the ongoing utility of existing bait sets - giving objective criteria for when and by what strategies baits should be updated. We also provide a method for quantifying and comparing antimicrobial resistance genes (ARGs) similar to what is used for RNAseq experiments. This approach improves comparison of ARGs across environments. Thus, the work provides an improved foundation for ARG future studies, while cutting across traditional areas of microbiology and extending beyond.

## Introduction

The World Health Organization (WHO) has cautioned of a “post-antimicrobial era”, in which worldwide deaths from antimicrobial resistance (AMR) could exceed 10 million each year by 2050 (Alsan et al., 2018). However, the U.S. Centers for Disease Control and Prevention (CDC) states that the post-antimicrobial era is already here, with 2.8 million AMR infections annually at a cost of $2 billion USD per year (O’Neil, 2014; CDC, 2019). These infections occur because microbes have developed the capacity to overcome microbicidal and microbiostatic activities of drugs. Resistance in bacterial populations is contingent on the characteristics of microbes (i.e., transmission mode, colonization, and pathogenicity) and the genetic basis for resistance.

For bacteria, resistance may be intrinsic, acquired, or may be due to point mutations conferring target modification (Sommer et al., 2010). Furthermore, plasmids, commonly found in Gram-negative bacteria, enable resistance genes to spread readily between bacteria and this spread poses a critical threat to human health (Buckner et al., 2018). Many drug-resistant bacteria have pathogenicity islands, which play a fundamental role in their virulence in humans (Schmidt and Hensel, 2004; Pendleton et al., 2013). Accurately identifying the resistome (i.e., the collection of AMR genes in a community) of environmental settings is vital for public health, as the spread of resistance and rise of new resistance elements are key public health concerns. AMR elements have been detected in a wide variety of reservoirs, including aquatic, terrestrial, food- and host-associated microbial communities (Oladeinde et al., 2018; Guitor et al., 2019; Mahnert et al., 2019; Oladeinde et al., 2019; Thomas et al., 2020). Increasing concerns over the AMR crisis have produced the need for AMR surveillance programs (e.g., at the United States Centers for Disease Control and Prevention). However, there is a lack of AMR surveillance programs for environmental reservoirs (Noyes et al., 2017; Arango-Argoty et al., 2018). Furthermore, existing methods to determine the resistome of complex environmental samples are not sufficient, and rely on laborious culture-based techniques, biased molecular-based methods, or extensive deep sequencing (Koser et al., 2014; Paterson et al., 2014; Van Camp et al., 2020). Furthermore, it is estimated that 99% or more of environmental bacteria cannot be readily cultured, leading to large bias in culture-based techniques (Luby et al., 2016).

Hybridization sequence capture (also known as capture enrichment sequencing, sequence capture, solution hybrid selection, target capture, or targeted sequence capture) is an enrichment technique that increases the proportion of DNA fragments of interest within DNA libraries (Mamanova et al., 2010; Teer et al., 2010). This method uses a set of DNA or RNA baits that are complementary to DNA sequences of interest to increase the proportion of target DNA fragments, and then characterize the DNA by MPS (Lasa et al., 2019). Hybridization sequence capture can be used to study AMR by designing baits to target the resistome. Hybridization sequence capture has several advantages, as limited *a priori* knowledge is required, there is no culture-bias, and it does not require extensive deep sequencing that is necessary to identify resistance elements in shotgun metagenomic libraries. In addition, the baits used to capture genes of interest can be designed to tolerate substantial sequence divergence (i.e., to detect new variants and overcome some level of genetic bias) while being highly sensitive. Hybridization sequence capture bait sets have been designed and used in microbial diversity studies by designing baits to the 16S rRNA (Gasc and Peyret, 2018; Beaudry et al., 2021), to study a defined set of pathogens or genes (e.g., virulence genes in *Vibrio* spp., human infecting viruses (Lasa et al., 2019; Metsky et al., 2019)), and to study antibiotic resistance (Noyes et al., 2017; Lanza et al., 2018; Guitor et al., 2019).

Prior to large-scale use of a novel bait set, effectiveness of the baits should be determined both *in silico* and *in vitro*. Of the previously designed bait sets for antimicrobial resistance, none have provided an *in silico* analysis (Noyes et al., 2017; Lanza et al., 2018; Guitor et al., 2019). *In silico* analyses are important because they characterize the efficiency of the bait set under ideal conditions and allow for easier comparison between published bait sets if similar bioinformatic methods are used. Moreover, *in vitro* validation of bait sets has been limited. Mock communities are often used to establish ground truth in microbial diversity studies; however, at the time of this writing no commercially available mock resistance community exists (Costea et al., 2017; Rausch et al., 2019). As such, several previous studies have limited their validation to host-associated samples (i.e., gut or stool samples) (Noyes et al., 2017; Lanza et al., 2018; Guitor et al., 2019) or phenotypic data, with only one study making a simple mock metagenome (Guitor et al., 2019).

The use of hybridization bait capture for antimicrobial resistance or other rapidly evolving genes of interest poses difficult challenges. Antimicrobial resistance genes are constantly evolving and being identified; for example between CARDv1.0.0 and CARDv3.1.1 a total of 676 resistance genes have been added (Alcock et al., 2020). If genes are not present in the reference database when baits are designed, then it is difficult to know if the bait set can capture new genes that are subsequently added to the database. This problem is similar to that of the Greek god Sisyphus, as soon as bait set design is complete then new ARGs will soon be added to the reference database(s) and design must start again. Similar to Sisyphus’s boulder rolling down the hill again, bait redesign is a never-ending chore. Thus, we examine how and when hybridization capture baits for the resistome should be updated, providing a guide to how researchers using hybridization capture can escape the fate of Sisyphus.

Here, we examine a targeted ARG sequence capture method (i.e., AMR-cap) to enrich metagenomic shotgun libraries for antibiotic resistance genes. Our bait set was designed from all genes in the CARD v1.1.2 and PAIDB v2.0 (Yoon et al., 2015; Alcock et al., 2020), which are reference databases for bacterial resistance genes and pathogenicity islands. We test our bait set *in vitro* on mock communities and diverse environmental samples, and provide *in silico* comparisons to previously designed hybridization capture bait sets (Guitor et al., 2019). Lastly, we provide evaluation methods for when to update hybridization capture bait sets, as the rapid identification of new ARGs poses a unique challenge to resistome bait sets.

## Results

### Design of Resistance Baits

Sequence entries in the Pathogenicity Island Database (PAIDB) v2.0 and CARD v1.1.2 were combined into a master reference source comprising roughly 19.6 Mbp of sequence space across 3,565 unique sequences (Yoon et al., 2015; Alcock et al., 2020). We first tiled 120nt baits approximately end-to-end across all these sequences (166,325 bait candidates), and then compressed these using USEARCH (Edgar, 2010) by eliminating any baits that were at least 90% identical with a minimum of 108 bp overlap with another bait (111,546 remaining). To select a subset of these baits that could serve to broadly sample the reference sequence space, we used MegaBLAST (Morgulis et al., 2008) to measure how many target reference intervals each bait might match. Starting with the bait with the highest number of interval hits (i.e., the most ‘ubiquitous’ bait) from the 111,325 survivors, we removed from consideration any other baits that also covered any of the same sequence space as that bait and retained the next most-ubiquitous bait among the survivors. This process was repeated until reaching roughly 10,000 bait candidates (part 1), which is anticipated to cover roughly 9.4 Mbp of the initial target space. Among the remaining candidates, another ~8,400 baits that only matched one location in the remaining target reference space were retained, keeping up to 10 baits per reference entry (part 2). We repeated the procedure used to generate part 1, but for the reference sequence intervals still un-baited to reach ~20,000 baits (part 3). This combination of two ‘general-purpose’ sets (parts 1 and 3) and one ‘specific’ set (part 2) comprises 19,933 total bait sequences, which is anticipated to cover roughly 11.0 Mbp of the original 19.6 Mbp target space.

### Sequencing Summary Statistics

For the unenriched metagenomic data, the aggregate number of total raw read pairs ranged from 4,773,192 in the resistance mock community to 37,789,368 wastewater treatment plant samples (Table 1). The highest percentage of reads retained after processing in the *de novo* assembly approach (see *Data Processing and Analysis*) was 82% in AMR-cap enriched poultry litter samples, and the lowest was 13% in unenriched built environment samples (Table 1). The percentage of reads retained after processing was greater than 30% in all other sample and library types.

**Table 1:**
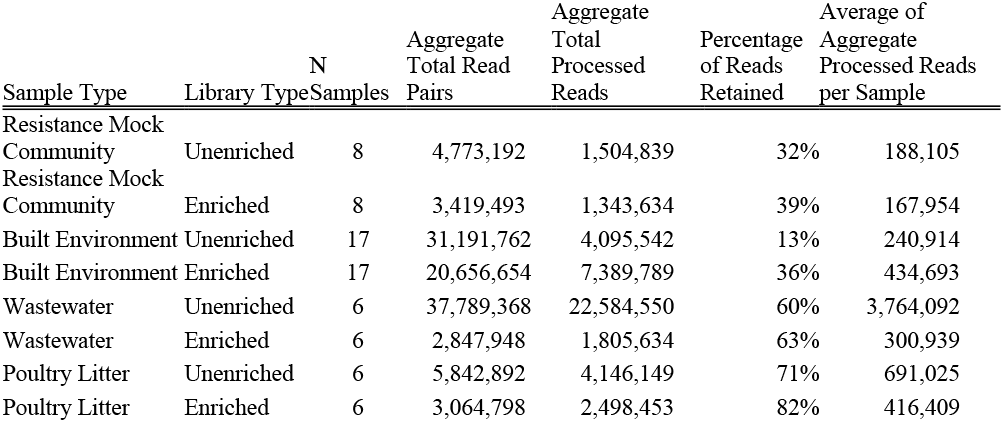
Summary of total Illumina HiSeq PE150 sequences available for each sample type using de-novo assembly with the AMRFinder database.

For the reference-based assembly, the aggregate number of high-quality reads by sample type in unenriched libraries ranged from 759,615 in mock community samples to 10,868,785 in wastewater samples (Table 2). The percent reads mapping to our combined target file in unenriched libraries ranged from 0.23% in the built environment samples to 4.72% in the resistance mock community samples (Table 2). In the reference-based assembly for AMR-cap enriched libraries, the number of high-quality reads by sample type ranged from 685,531 in wastewater samples to 1,926,747 in built environment samples (Table 2). Processed reads were mapped to our combined target file and ranged from 31.93% mapping in built environment samples to 84.80% mapping in the resistance mock community samples (Table 2).

**Table 2:** Summary of total sequences, percent mapping, fold change, and empirically achievable enrichment targets for reference-based assembly approach with averages for each sample type.

| Sample Type | Unenriched<br>No. of High<br>Quality Reads | Unenriched<br>PE150 Total<br>Mapped | Unenriched<br>Percent<br>Mapped | Enriched No.<br>of High<br>Quality Reads | Enriched-PE150<br>Total<br>Mapped | Enriched<br>Percent<br>Mapped | Fold<br>Change | 90% on<br>target<br>empirically<br>achievable |
| --- | --- | --- | --- | --- | --- | --- | --- | --- |
| Resistance<br>Mock<br>Community | 759,615 | 35,970 | 4.72% | 858,344 | 729,721 | 84.80% | 17 | 19 |
| Built<br>Environment | 1,244,146 | 3,856 | 0.23% | 1,926,747 | 645,221 | 31.93% | 136 | 387 |
| Wastewater | 10,868,785 | 87,361 | 0.79% | 685,531 | 315,396 | 45.80% | 57 | 114 |
| Poultry Litter | 1,692,429 | 4,438 | 0.27% | 853,560 | 586,455 | 67.80% | 248 | 331 |

Hybridization capture enrichment efficiency was estimated by calculating the fold-change (increase) for each sample and library type, and an average was obtained for each sample type (Table 2). The lowest fold change was observed in the resistance mock community samples at 17. It should be noted, however, that if the sample had been enriched to 90% on target reads (i.e., about the maximum achieved in hybridization capture) the fold change would be 19 (Table 2). The environmental samples fold change ranged from 57 in the wastewater samples to 248 in the poultry litter samples, with fold change >100 being achievable in all environmental sample types when 90% of reads are on target (Table 2).

### In silico Simulation

We created two simulated data sets for *in silico* capture experiments of our baits and those of Guitor et al. (2019), all of which achieved ≥99.99% mapping of the expected number of sequences (Table 3). We observed a higher percentage of total mapped reads in our *in silico* resistance mock community than our *in vitro* resistance mock community (*cf*. Tables 2, 3). For example, the average total mapping of the simulated data is 100% in the resistance mock community (Table 3) whereas it was 84.80% in the empirical data (Table 2).

**Table 3:**
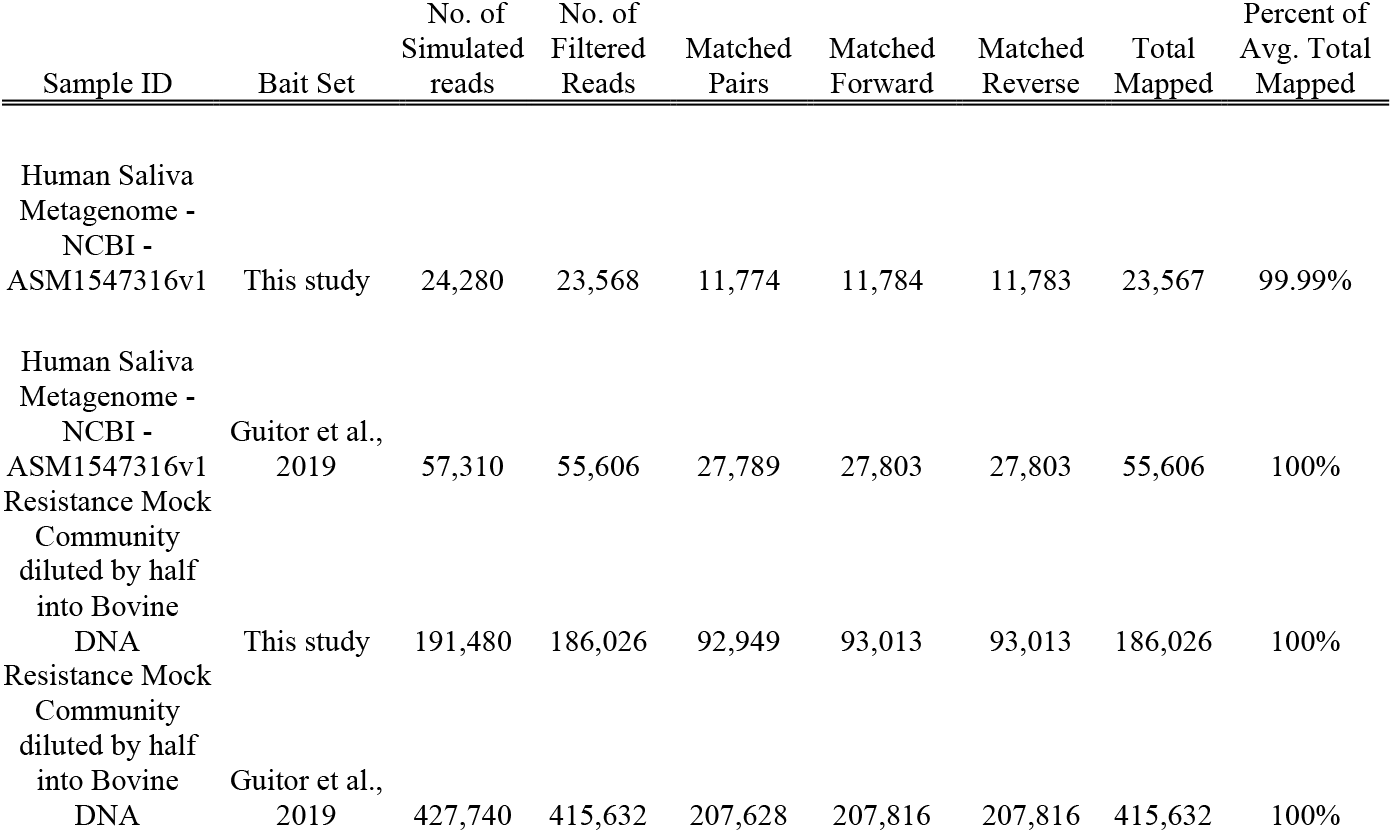
Summary of simulated Illumina PE150 data for a human saliva metagenome and the mock resistance community developed in our study.

### Validation in Resistance Mock Community using a De Novo Assembly Approach

We initially prepared unenriched metagenomic libraries and enriched AMR-cap libraries on a resistance mock community containing 203 different ARGs (Supplemental Materials Table 1). Overall, using *de novo* analysis, we detected 145 different ARGs in our AMR-cap enriched library representing every resistance class present in the original mock community (Supplemental Figure 3). In our enriched libraries (Figure 1, panel A, column 2), we were able to detect all expected ARGs within five resistance classes, (blocks indicated with 100% and bright green), more than 66-88% of expected ARGs in four more resistance classes, and only two resistance classes performed poorly (beta-lactam - 20%; colistin – 14%). In contrast, in our unenriched library (Figure 1, panel A, column 3), most resistance classes performed poorly; we were unable to detect any ARGs in two classes of resistance: colistin and phenicol/quinolone. A Wilcoxon rank sum test revealed a significant difference in the number of ARGs identified in AMR-cap enriched libraries compared to unenriched metagenomic libraries (p-value = 0.0033) (Supplemental Figure 2).

**Figure 1:**
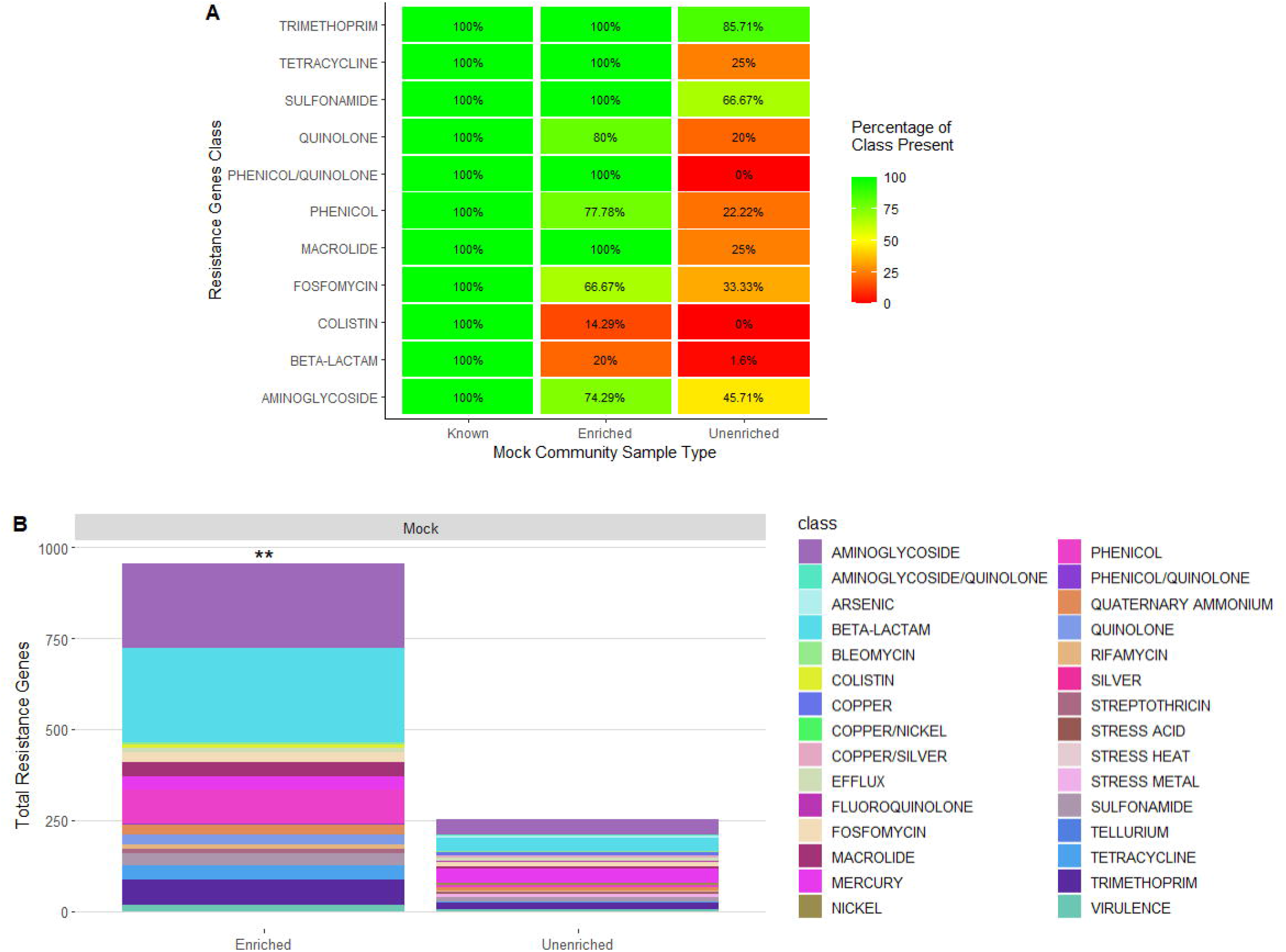
(A) Aggregate equalized data heat plot of the known content of the mock community broken down by resistance class on the y-axis. Library type and known content are labeled on the x-axis. Percentage of class present ranges from 0% (red) to 100% (green). (B) Aggregate equalized data for total resistance genes (y-axis) found in the AMR-cap enriched and unenriched libraries (x-axis) broken down by resistance class. Stars represent a significant difference (p-value ≤ 0.05) between AMR-cap enriched and unenriched libraries.

### Validation in Environmental Samples using a De Novo Assembly Approach

In this series of analyses, it was critical to down-sample sequences so that all samples had identical numbers of sequences (Supplemental Figures 4 & 5) (See *Data Processing and Analysis).*

Wilcoxon rank sum test was used to compare total ARGs identified in unenriched and AMR-cap enriched libraries by sample type (i.e., wastewater, poultry, and built environment) in our data analyzed with a *de novo* assembly approach (Supplemental Figure 2). Our analysis revealed that there were significant differences (p-value ≤ 0.05) between the unenriched and AMR-cap enriched libraries for all sample types (Figure 2, Supplemental Figure 2).

**Figure 2:**
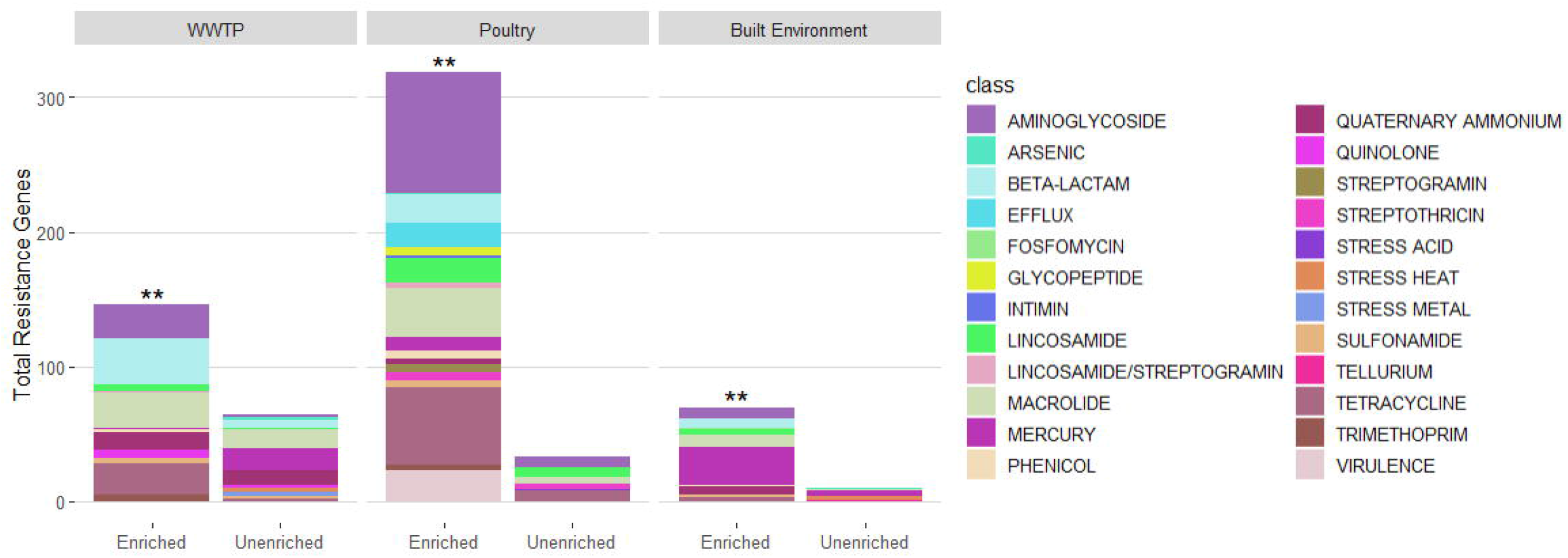
Aggregate equalized data for stacked bar plot of total resistance genes (y-axis) by sample top (top) found in AMR-cap enriched and unenriched libraries (x-axis) broken down by resistance class. Stars represent a significant difference between AMR-cap enriched and unenriched library types.

In the unenriched built environment samples, a total of 10 resistance genes were identified, of which eight were unique (i.e., only found once) and belonged to five classes of resistance (Figure 2, Supplemental Figure 1 & 3). In comparison, AMR-cap enriched libraries were able to identify 70 total resistance genes of which 36 were unique resistance genes corresponding to 10 different classes of resistance (Figure 2, Supplemental Figures 1 & 3). On average, we detected 8.57 resistance genes per AMR-cap enriched built environment samples and 1.67 resistance genes for unenriched built environment samples (p-value = 0.0058) (Supplemental Figure 2).

In samples obtained from wastewater treatment plants, AMR-cap was able to identify 12 classes of resistance and 146 total resistance genes, with 54 being unique resistance genes. In comparison, in the unenriched libraries, we were able to identify 11 classes of resistance and 64 total resistance genes, of which 27 were unique (Figure 2, Supplemental Figures 1 & 3). Furthermore, on average, in wastewater treatment plant samples, we were able to identify 28.8 resistance genes per enriched sample and 9.83 resistance genes per unenriched sample (p-value = 0.0078) (Supplemental Figure 2).

Analysis on poultry litter samples revealed that AMR-cap identified 18 classes of resistance and 318 total resistance genes, of which 78 are unique. For the unenriched libraries, 6 different classes of resistance and 33 total resistance genes were identified, of which 16 are unique (p-value = 0.0079) (Figure 2, Supplemental Figures 1 & 3).

### Reads Per Kilobase per Million mapped reads (RPKM) Analysis

In this series of analyses, it is possible to use all available sequencing reads (see *Data Processing and Analysis*) (Beaudry et al., 2021). An RPKM analysis of our resistance mock community revealed that resistance genes in enriched libraries are sequenced at a much higher sequencing depth than those in unenriched libraries, in all but one resistance class. The exception is fosfomycin, which only had one resistance gene (Figure 3). The RPKM analysis in the resistance mock community highlights the ability of enrichment to sequence classes of genes that are missed in the unenriched libraries (i.e., colistin). Furthermore, across enriched replicates of the resistance mock community, we observed particular resistance genes being sequenced deeply (i.e., red lines in Figure 3).

**Figure 3:**
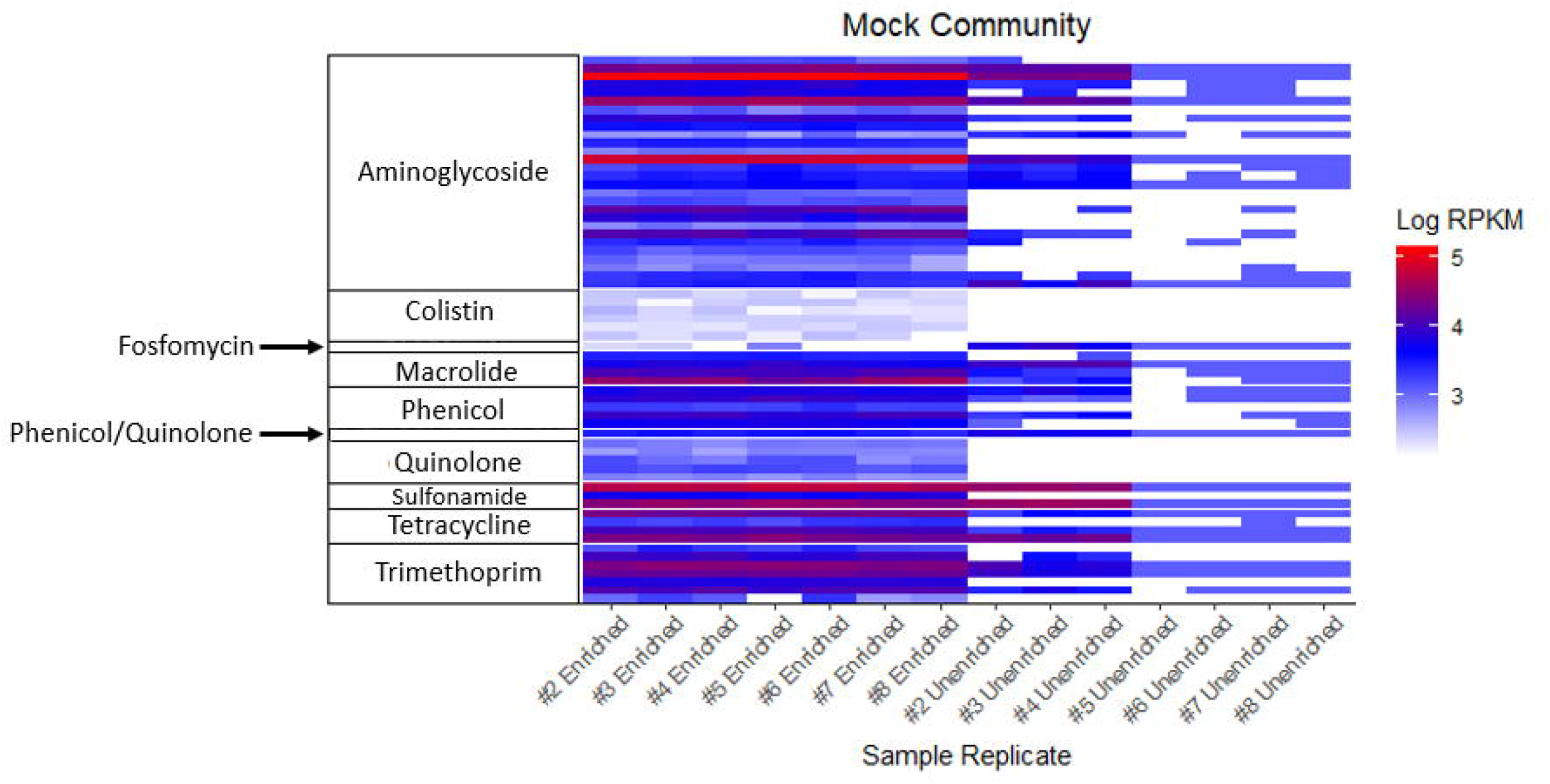
RPKM analysis of the non-equalized resistance mock community. Sample replicates are indicated on the x-axis with enriched samples on the left and unenriched samples on the right. RPKM Values are log transformed, a key is located on the right. Prominent classes are labeled on the left side.

The RPKM analysis of our three environmental sample types (i.e., poultry litter, wastewater, and built environment) shows the entire spectrum of results that AMR-cap enrichment can achieve (Figure 4). In built environment samples, we observe an increased sequencing depth across all resistance genes in AMR-cap enriched libraries compared to unenriched libraries. Furthermore, the RPKM analysis of built environment samples reflects the inability of unenriched metagenomic libraries to identify a broad range of resistance genes in the majority of these samples. In the poultry litter samples, enrichment increases the depth of sequencing for most, but not all resistance genes. Likewise, enrichment is shown to be highly effective on some samples (e.g., poultry litter sample 5). Thus, the poultry litter samples demonstrate that enrichment is valuable, but obtaining data from unenriched libraries also adds value for samples with relatively high numbers of ARGs. In the wastewater, however, the RPKM results demonstrate that deep sequencing of unenriched samples can recover more depth from a broad array of resistance genes versus more limited sequencing of enriched samples. Even in the wastewater samples, however, there are still a few ARGs detected in the enriched but not unenriched libraries.

**Figure 4:**
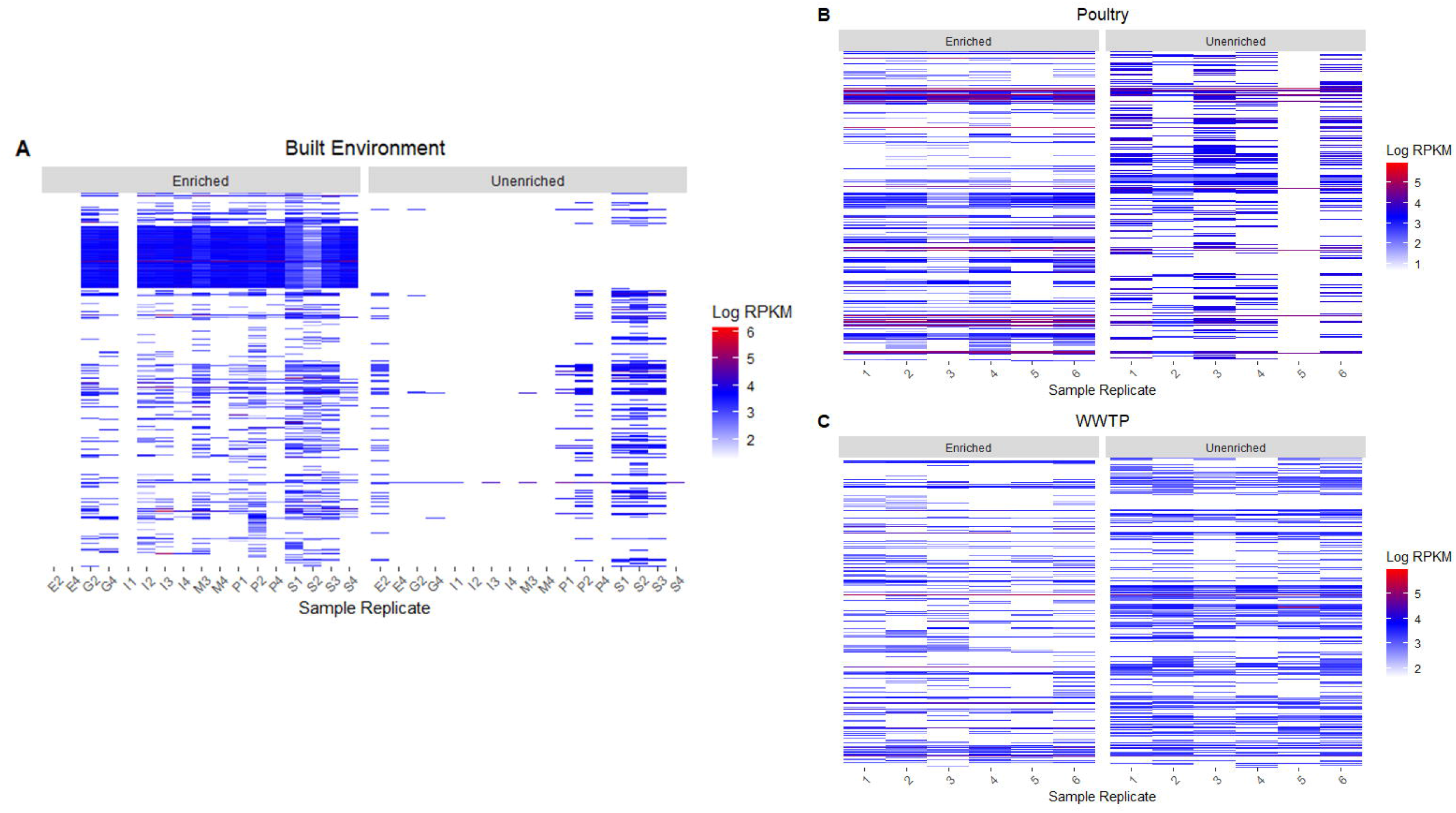
RPKM analysis of non-equalized environmental samples broken down by sample type: A) Built Environment, B) Poultry, and C) Wastewater Treatment Plant. Sample replicate is on the x-axis with enriched samples on the left and unenriched samples on the right. RPKM values are log transformed, a key is located on the right for each figure.

### In silico Bait Set and CARD Comparisons

A comparison between our bait set, the Guitor et al. (2019) bait set, CARDv1.0. and CARDv3.1.1 was performed as previously described (see Data Analysis). These comparisons were chosen for several key reasons: 1) at the time of analysis CARDv3.1.1 was the newest version of CARD, 2) pathogenicity islands are included in CARDv3, and 3) PAIDB is no longer available. Both bait sets were compared to two versions of CARD (CARDv1.0 and CARDv3.1.1). We found that our bait set, AMR-cap, was able to capture 2092 (99.1%) of 2110 ARGs in CARDv1.0, and the Guitor et al. (2019) bait set was able to capture 2083 (98.7%) of 2110 ARGs in CARDv1.0. Additionally, a comparison to CARDv3.1.1 was made, which is comprised of 2786 unique ARGs. Our bait set was able to capture 2503 (89.8%) ARGs, whereas the Guitor et al. (2019) bait set was able to capture 2351 (84.3%) ARGs (Figure 5). ARGs the bait sets did not capture are in the supplemental materials (Supplemental Tables 2 & 3).

**Figure 5:**
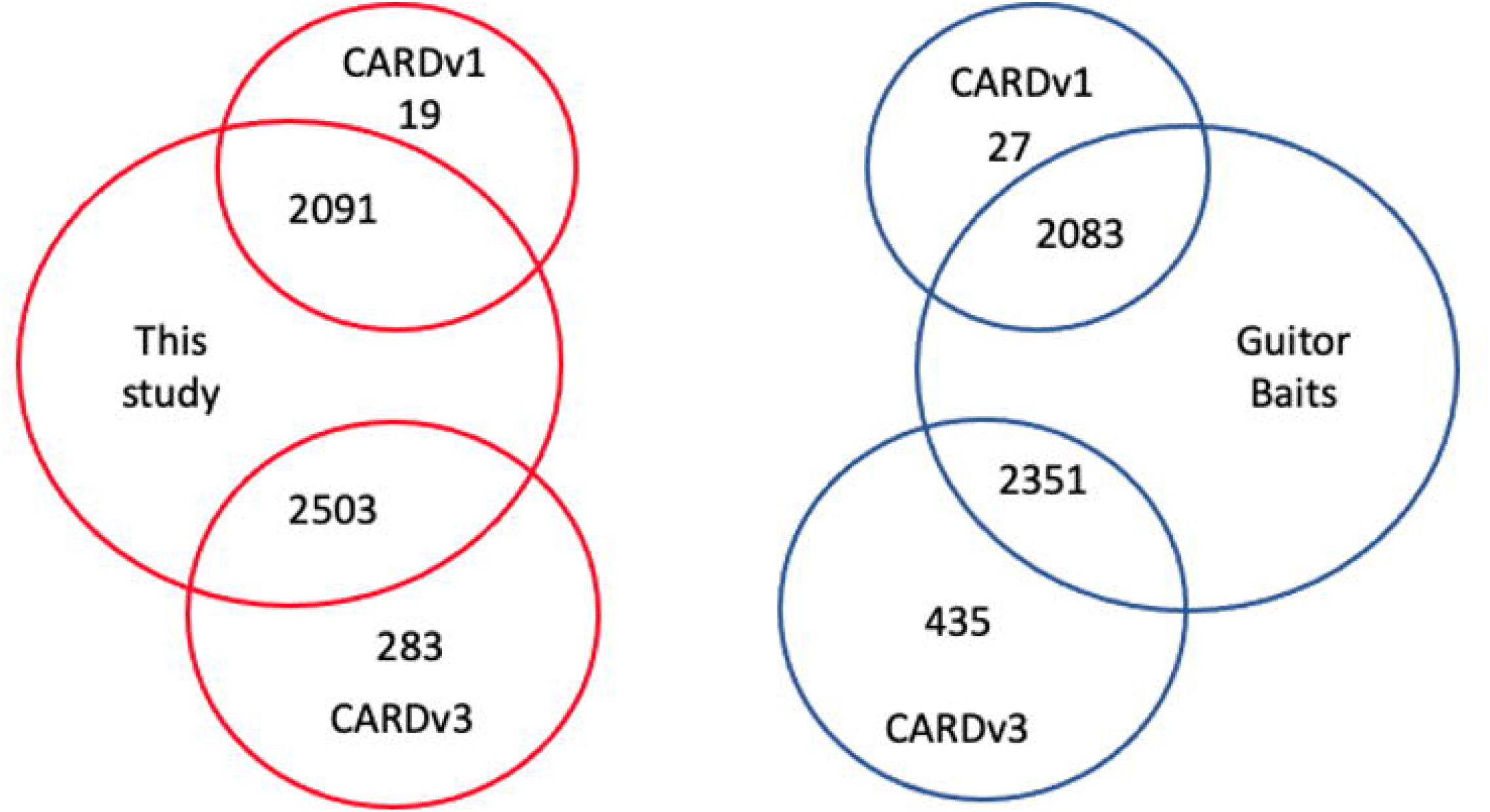
Venn diagram of the results of AMR-cap baits (red) and Guitor baits (blue) capturing genes in CARDv1 (top) and CARDv3 (bottom).

## Discussion

Given the limitations of currently available culture-dependent and culture-independent methods to study antimicrobial resistance in environmental samples, we sought to provide an alternative method to identify resistance genes in environmental samples by creating an antibiotic resistance hybridization capture assay (i.e., AMR-cap). Our study revealed several important findings: (1) AMR-cap achieves a higher percentage of on-target reads than unenriched libraires regardless of sample type, (2) AMR-cap is an efficient and cost-effective way to obtain ARG sequences, and (3) AMR-cap is able to capture novel ARGs in CARDv3.1.1 that were unknown when our bait set was designed. Our study was not the first to apply hybridization bait capture to antibiotic resistance. However, our meticulous bait design combined with *in silico* and *in vivo* experiments using a custom-built resistance mock community and wide variety of environmental sample types (i.e., agriculture, water, and built environment) reflects the capability of our bait set for studying complex environmental samples. Furthermore, this is the first study to provide a published framework for updating resistome hybridization capture bait sets (see below).

Enriching for genes of interest is an important technique to study AMR. A multitude of past studies have relied on PCR-based assays which are sensitive but require significant *a priori* knowledge of ARGs of interest and do not provide the depth of information that can be acquired from unenriched metagenomic shotgun libraries. Other previous studies have used hybridization capture to enrich for ARGs (Noyes et al., 2017; Lanza et al., 2018; Guitor et al., 2019). Guitor et al. (2019) tested their hybridization capture bait set for antibacterial resistance on a mock community and found that ~70-80% of their enriched reads map to bait-targeted regions. Our bait set had similar results, with 85% of reads in our AMR-cap mock community enriched library mapping to bait-targeted regions, in comparison to the unenriched metagenomic library which had 4.2% of reads mapping to bait-targeted regions (Table 2). Using a *de novo* analysis approach, we find on average 99 ARGs per sample in enriched libraries from mock communities comparted to 38 ARGs per sample from unenriched metagenomic libraries (Supplemental Figure 2). More specifically, in our mock resistance community, we find that two classes of resistance (i.e., beta-lactam and colisitin) did not perform as well as other classes. This may be in part due to the design of the mock resistance community, which included 125 ARGs for beta-lactam, which is more than all other resistance classes combined (i.e., 78 ARGs) (Supplemental Table 1). Additionally, with the methods employed in this study we cannot tell all of the 125 beta-lactam sequences apart, even though they are likely present among the reads. Therefore, our AMR-cap assay achieves a very high percentage of on-target reads and increases the number of ARGs identified regardless of bioinformatic analysis method.

Environmental samples are particularly challenging to work with due to their complex nature. Validation of AMR hybridization capture bait sets in previous studies has been limited to host-associated samples, with only one study testing wastewater samples (Noyes et al., 2017). In our study we tested a variety of environmental samples including poultry litter, built environment samples, and wastewater treatment plant influent and effluent. When the number of reads was normalized between AMR-cap enriched and unenriched libraries for each sample type, we found that our AMR-cap assay identifies significantly more resistance genes and classes of resistance in all environmental sample types (Figure 2). Many of these ARGs come from clinically important groups including beta-lactam resistance, tetracycline resistance, and aminoglycoside resistance. More specifically, built environment and poultry litter AMR-cap enriched samples identified several clinically important genes that were not identified in unenriched libraries. This included blaOXA, which confers carbapenem resistance (i.e., an antibiotic of last resort) and are widespread in *Acinetobacter baumannii*, and *mecA*, which encodes an extra penicillin-binding protein and is one of the most impactful antimicrobial resistance genes to human health worldwide (Rolo et al., 2017). In addition, we detected blaTEM genes in AMR-cap enriched samples from built environments, but not in unenriched metagenomic libraries, thus highlighting the ability of our method to identify beta-lactam ARGs in environmental samples. Noyes et al., 2017 detected TEM beta-lactamases in 29 of 32 enriched fecal samples, but not in any unenriched samples. This indicates that these ARGs are low abundance within each sample and could not be detected without hybridization capture. A similar trend is observed in the non-normalized data (Supplementary Materials Figure 5). However, more metal resistance genes and classes of resistance were identified in unenriched wastewater treatment plant samples due to the high levels of metal resistance genes in wastewater (Supplemental Figure 5).

An additional benefit of our AMR-cap assay is that it increases the number of high-quality reads obtained from enriched libraries compared to unenriched libraries. This phenomenon is apparent in the built environment samples, where only 13% of reads from unenriched libraries were retained after quality filtering and 36% of reads were retained from the AMR-cap libraries (Table 1). The number of reads obtained in our unenriched libraries for low biomass samples from built environments is similar to other studies (Gibbons et al., 2015; Brooks et al., 2017; Chen et al., 2017).

Our RPKM analysis revealed several important findings. Foremost, we find that AMR-cap enrichment leads to varying degrees of success in different types of environmental samples (Figure 3, Figure 4). In the resistance mock community and built environment samples, AMR-cap enrichment is highly effective for sequencing more resistance genes at a greater depth. Noteworthily, without AMR-cap enrichment few resistance genes, if any, are sequenced in the unenriched libraries from built environments. Those that are identified had a much lower sequencing depth than those in AMR-enriched libraries. As previously mentioned, samples obtained from built environments are often of low microbial load, and without enrichment resistance genes are below detection levels (Gibbons et al., 2015; Brooks et al., 2017; Chen et al., 2017). Conversely, we found that AMR-cap enrichment has the opposite effect on samples obtained from wastewater treatment plants. Wastewater samples are notorious for having a high microbial load and, in comparison, to other sample types have higher levels of ARGs (Amalfitano et al., 2018). The poultry litter samples represent a middle ground between the built environment and wastewater samples, as some samples and resistance genes are sequenced at deeper levels with AMR-cap enrichment. Thus, our AMR-cap enrichment may be best suited for samples with a low- to average microbial load where AMR-cap enrichment identifies ARGs with a greater sequencing depth and coverage than before.

During the 6 years between CARDv1.0.0 and CARDv3.1.1, 676 resistance genes were added to the database (Alcock et al., 2020). We tested our bait set and the bait set of Guitor et al. (2019) against CARDv1.1 and CARDv3.1.1, finding that our bait set was able to capture 152 more resistance genes than the Guitor bait set. This indicates that our AMR-cap bait set has a somewhat better ability to identify novel ARGs than the Guitor bait set. Our bait set was designed to maximize the probability of obtaining sequences similar to the baits (i.e., by using 120-mer baits), in comparison to the Guitor baits which focus more on specificity, thus using 80-mer baits. In our analyses, we assume that both bait sets will tolerate 15% nucleotide divergence from the reference sequence, but longer baits allow for more divergence which is key as many newly characterized ARGs only differ from other ARGs by a few nucleotides (Li et al., 2013; Guitor et al., 2019).

We can make extrapolations of expected future performance of AMR-cap based on previous updates to CARD. On average, the CARD database added ~110 resistance genes per year (Supplemental Table 4). Our bait set had an ~60% success rate at identifying new resistance genes added to CARDv3.1.1 vs. CARDv1.0, thus for every seven new resistance genes added to CARD our baits are expected to identify approximately four. Based on these calculations and assuming a similar pace of discovery in the future, we estimate that over a 10-year period (i.e., from CARDv1 to CARDv5) ~1100 new resistance genes will be added. If our bait set performs similarly in future iterations of CARD, then our bait set will be able to capture 85% of resistance genes in the CARDv5 database (Supplemental Table 4). A similar trend is observed after a 16-year period (i.e., to CARDv8), where our bait set can capture 80% of resistance genes in the database (Supplemental Table 4). When 15-20% of genes are not identified, it is likely best to extend the capability of AMR-cap through a supplemental set of baits that capture new clusters of genes (i.e., a patch). In contrast, over a 20-year period (i.e., from CARD v1 to CARDv10) ~2200 novel resistance genes are predicted to be added to CARD, and our bait set will only be able to capture ~67% of them (Supplemental Table 4). As multiple patches are increasingly difficult to design and each patch adds cost to the bait set, by the time CARDv10 is released, it is likely to be worthwhile to redesign the entire bait set. Thus, based on the current pace of CARD updates, we expect AMR-cap to require a major patch about once per decade and a redesign once per 20 years. However, this expected timeline is dependent upon most periodic additions to the database consisting of sequence variations including insertions or recombinations of previous ARGs; discovery and addition of a major new class of ARGs with major clinical public health impact would be cause for earlier additions or redesigns. Likewise, major global increases in sequencing capacity and thus identification of new ARGs may alter this timeframe. Nevertheless, minor patches of critical new ARGs, using traditionally synthesized individual biotinylated oligonucleotides can be added at any time.

We find that the AMR-cap assay requires far fewer reads than unenriched metagenomic libraries, consequently, allowing AMR-cap libraries to be characterized on Illumina MiSeq instruments at a reasonable cost. MiSeqs are widely available in public health laboratories across the world. Thus, this approach opens opportunities for many more labs to initiate such work. Our bait set provides a valuable tool for assessing antibacterial resistance in a variety of settings that require public health surveillance (e.g., water, clinical built environments, etc.) (Beaudry et al., 2021). Our ready-to-use publicly available bait set is a cost-effective way to monitor AMR (i.e., baits can be purchased from Arbor Biosciences as Design ID# D10091Rsst; or the baits can be synthesized by researchers using the bait sequences), as other studies have shown the cost of enrichment with purchased kits can be as low as $11.72 per sample (Glenn and Faircloth, 2016; Beaudry et al., 2021). However, further research is needed to fully utilize this tool for AMR surveillance including understanding risk in the context of genomic signal and defining the criteria for genomic data on AMR needed for a public health response (Port et al., 2014; Costea et al., 2017; Noyes et al., 2017). Lastly, a comparison of traditional surveillance approaches to the genomic method outlined in this paper is necessary.

This study had several limitations. The bioinformatic pipelines used for *de novo*, reference-based, and RPKM analysis produced differing results. This may be due to the difference in reference databases (e.g., AMRfinder vs CARD), variation in trimming/quality filtering programs (e.g., trimmomatic vs BBDuk), or quality filtering parameters (e.g., minimum length, q-score). Future work should systematically investigate the effects of these and other analysis options and provide easily repeatable pipelines for AMR-cap analyses, so results from different studies are readily comparable. Furthermore, future work could test varying hybridization times, sequencing depths, and sequencing read length to have a greater understanding of how this may affect the detection of different resistance classes (e.g., beta-lactams). In addition, we do not provide a direct *in vitro* comparison of other resistance bait sets to our AMR-cap assay. Future research may benefit from such a comparison.

In conclusion, our data demonstrate that our AMR-cap assay targeting >3500 sequences in CARD efficiently enriches for ARGs in resistance mock communities and a variety of complex environmental samples. Specifically, in complex environmental samples we achieved >200-fold enrichment, thus reducing the need for extensive deep sequencing to identify ARGs allowing for increased flexibility on sequencer choice. We believe that our bait set is a valuable and cost-effective tool for public health surveillance of AMR in key areas of concern (i.e., water, agriculture, hospitals). Furthermore, we provide an *in silico* comparison of our bait set to previously designed resistome bait sets and demonstrate the adaptability of our bait set to capture novel ARGs.

## Experimental Procedures

### Resistance Mock Community Design and Testing

A mock resistance community was created by extracting DNA from AMR isolates provided by the CDC & FDA Antibiotic Resistance Isolate Bank (AR Bank) (https://www.cdc.gov/drugresistance/resistance-bank/index.html). A mock community containing 49 isolates (Supplemental Table 1) was produced. These isolates belong to different AR Bank resistance panels (i.e., Enterobacteriaceae Carbapenem Breakpoint Panel, Gram Negative Carbapenemase Detection Panel, Isolates with New or Novel Antibiotic Resistance, and Ceftolozane/tazobactam Panel), and in total include eleven bacterial species (i.e., *Enterobacter cloacae, Acinetobacter baumannii, Klebsiella pneumoniae*, *K. ozaenae*, *K. oxytoca*, *Pseudomonas aeruginosa*, *Morganella morganii*, *Proteus mirabilis*, *Enterobacter aerogenes*, *Escherichia coli*, and *Serratia marcescens*). Each isolate was added in equal proportion to our mock community.

### Samples and DNA Extraction

Samples included in the study were collected from a variety of environmental sources (for N values see Table 1). Built environment samples were collected using sponges and swabs from healthcare facilities, offices, and gyms throughout the United States. Following collection, DNA samples were extracted with the PowerSoil DNA Isolation kit (Qiagen, Hilden, Germany). Wastewater treatment plant (WWTP) influent and effluent were collected by grab sampling, and DNA was extracted using previously described methods (Teachey et al., 2019). Poultry samples were collected from poultry litter, DNA was extracted and purified using previously described methods (Oladeinde et al., 2019). Mock community isolates were extracted using the PowerSoil DNA Isolation kit (Qiagen, Hilden, Germany).

### Metagenomic Libraries

Metagenomic shotgun libraries were prepared using New England BioLabs Ultra FS II library kit (New England BioLabs, Ipswich, MA) and cleaned with a 1:1 ratio of Sera-Mag speedbeads after ligation (Rohland and Reich, 2012). Following ligation, samples were tagged with iTru primers (Glenn et al., 2019) and quantified with a Qubit 2.0 Fluorometer DNA high sensitivity assay kit (Thermofisher, Waltham, MA). Samples were then cleaned with a 1:1 ratio of speedbeads and pooled into groups of 8-12 for enrichment or set aside for sequencing. A positive control of 2.5 ng/μL of *E. coli* and a negative control of molecular grade water were included.

### Resistome Capture Enrichments

Pooled libraries were then used to perform hybridization capture with biotinylated baits using a custom Arbor Biosciences myBaits kit (Arbor Biosciences, Ann Arbor, MI). The kit was used following manufacturer’s protocol (v3-5.01) with 16-18-hour hybridization at 65°C for all samples. Following hybridization, Dynabeads M-280 Streptavidin magnetic beads (Life Technologies, Carlsbad, CA) were used for enrichment. A post-enrichment amplification was performed using Illumina P5/P7 primers and KAPA HiFi HotStart reagents. The cycling conditions were as follows: 98°C for 45s, followed by 16-28 cycles of 98°C for 20s, 60°C for 30s, and 72°C for 60s, and then a final extension of 72°C for five minutes. P5/P7 PCR consisted of 25 cycles for the resistance mock resistance community, 25 cycles for the built environmental samples, 18 for the poultry litter samples, and 18 cycles for the WWTP samples. PCR product was cleaned with Sera-Mag beads, quantified on a Qubit 2.0 fluorometer DNA high sensitivity assay kit (Thermofisher, Waltham, MA) and pooled in equimolar ratio for sequencing.

### Simulating Resistome Target Enrichment Data

To test the efficiency of our bait set and Guitor et al., 2019 bait set under ideal conditions, we did an *in silico* analysis to determine how well our bait sets work during an error- and bias-free hybridization process (Alcock et al., 2020). Two metagenomes were used, one a human saliva metagenome obtained from NCBI (ASM1547316v1) and one custom-built metagenome to match our mock resistance community previously described. Fasta files containing sequences for each of the bait sets were mapped to each of the metagenome fasta files using the Burrows-Wheeler Aligner (bwa) v0.7.17, simulating an error-free and bias-free hybridization process (Li and Durbin, 2009). The SAM files created by the mapping process were converted into BAM files using SAMtools v0.1.19 (Li et al., 2009). Then the mapping coordinates obtained from each of the bait sets on the reference metagenomes were used to extract the sequences +/− 350bp upstream and downstream of the coordinate’s position, wherever possible. This was done to simulate the hybridization of the bait to the core of an ~800 bp fragment while also acquiring the flanking regions typically captured when using biotinylated baits. Paired-end 150 bp fastq reads were simulated at 10X coverage using the software ART 2016.06.05 and the reference sequences from each metagenome that were created with each bait set (Huang et al., 2012).

The *de novo* and reference-based assembly were performed as described below (see Data Processing and Analysis). The number of ARGs present in each metagenome with each bait set was recorded. Additionally, for the reference-based assembly the human saliva metagenome and the resistance mock community were used as a reference for the reads simulated with each bait set. The number of reads that mapped to the reference metagenomes were recorded (Table 3).

### Sequencing

Resistome enriched libraries and unenriched metagenomic libraries were sequenced on an Illumina HiSeq X Ten using a HiSeq X Ten Reagent Kit v2.5 (300 cycles) (Illumina, San Diego, CA) to obtain PE150 reads. Data was demultiplexed using bcl2fastq2 v.20 (Illumina, San Diego, CA).

### Data Processing and Analysis

For *de novo* assembly, Illumina adapters were removed using Trimmomatic v0.39 using a lead and trailing quality of three Phred-33 (Bolger et al., 2014). To equalize the number of reads found across sample types, 500,000 reads were randomly selected (i.e., subset of the data) from mock community, wastewater, and poultry samples. One million reads were randomly selected from built environment samples. Random selection was done using Seqtk (Shen et al., 2016). Samples not equalized by the number of reads are included in the supplemental materials (Supplemental Figures 4-5). Samples were quality filtered with a sliding window of 4:15 and minimum length of 100. SPAdes v3.14.1was used to assemble reads into contigs using the options -meta and -plasmid (Nurk et al., 2017). To query contigs, AMRFinderv3.9.8 with the option nucleotide, threshold of 0.5 (see Supplemental Figure 6 for information on thresholds), and plus were used to determine the number of hits for each gene in each resistance category. The number of hits for each resistance category and antimicrobial resistance gene were recorded.

For reference-based assembly, total read data of all sequences were downloaded into Geneious Prime v2020.2 or v2021.1 and paired. BBDuk was used to trim Illumina TruSeq adaptors, with a kmer length of 21 and trim partial adapters from ends with a kmer length of 8. Additionally, reads smaller than 50 bp were discarded and a minimum quality score of 25 was used. Our targets file (i.e., CARDv1.1.2 and PAIDBv2.0) was uploaded into Geneious and used as a reference to map sequence reads using BBMap version 38.84 with default settings.

To query bait sets (i.e., Guitor et al., 2019 and our bait set) and databases (i.e., CARDv1.0.0 and CARDv3.1.1) against each other, BWA v0.7.17 with the option mem and default settings was run (Li and Durbin, 2009). The baits were used as a reference to align genes against the baits.

To determine the abundance of ARGS as expressed in RPKMs (reads per kilo base per million mapped reads), total read data were trimmed and trimmed paired reads were mapped to CARDv3.1.1 using BWA mem 0.7.17 (Li and Durbin, 2009). A custom bash script was used to subset mapped reads from the SAM alignment file and calculate the length of each gene from CARDv3.1.1 database. RPKM values were calculated using the following formula: RPKM = numReads/(geneLength/1000 *totalNumberReads/1,000,000). Genes with less than 10 mapped reads were discarded. Classes and subclasses were matched to genes from using CARDv3.1.1 and AMRfinderv3.9.8.

R statistical software (R Development Core Team, 2010) was used to make plots and run statistical analyses. The Shapiro-Wilk normality test was used and indicated that de-novo data were not normally distributed. Wilcoxon Rank-Sum tests were used to compare the median number of ARGs identified in the enriched and unenriched libraries between each sample source. Data processing, summarization, and analysis was completed using base R and the ‘tidyverse’ program library (Wickham et al., 2019). Data visualizations were created using the ‘ggplot2’ library.

## Data Availability Statement

The data is publicly available through the NCBI Sequence Read Archive (SRA) BioProject ID PRJNA746311.

## Author Contributions

TG conceived the project. MB, JT, TK, NV, AS, LR, AP, and AO designed the experiments. MB performed the experiments. JT, AD, and JE designed the baits. MB, RB, and AS analyzed the data. MB and WN made the figures. SS, TG, and OR provided the funding. MB wrote the manuscript. All authors critically reviewed, edited, and approve of this work.

## Disclaimer

The findings, opinions, and conclusions in this report are those of the authors and do not necessarily represent the views, official positions or policies of the Centers for Disease Control and Prevention (CDC), U.S. Department of Energy (DOE), the U.S. Department of Health and Human Services (HHS), the Public Health Service (PHS), or the U.S. Department of Agriculture (USDA). Use of trade names is for identification only and does not imply endorsement or recommendation for use by the CDC, DOE, HHS, PHS, or USDA.

## Acknowledgments

We thank: Marissa Howard, Paula Bartlett, Dale Green, Elizabeth Ottesen, and Jason Westrich for their generous contributions to this work.

