## Supplemental Materials for "Escaping the Fate of Sisyphus: Assessing Resistome Hybridization Baits for Antimicrobial Resistance Gene Capture"

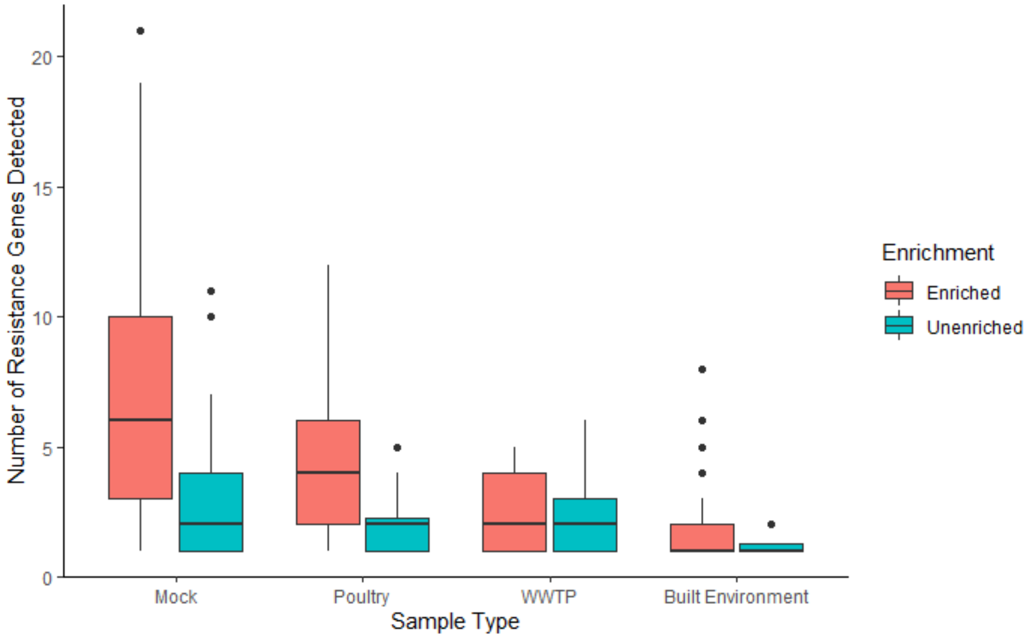


Supplemental Figure 1 Box and whisker plot of the distribution of gene occurrence broken down sample type (x-axis) and library type (red – enriched, teal – unenriched). The 50th percentile is represented by a line in the middle of the box, the 25th and 75th percentiles are the edges of the box. Outliers are represented by stars.

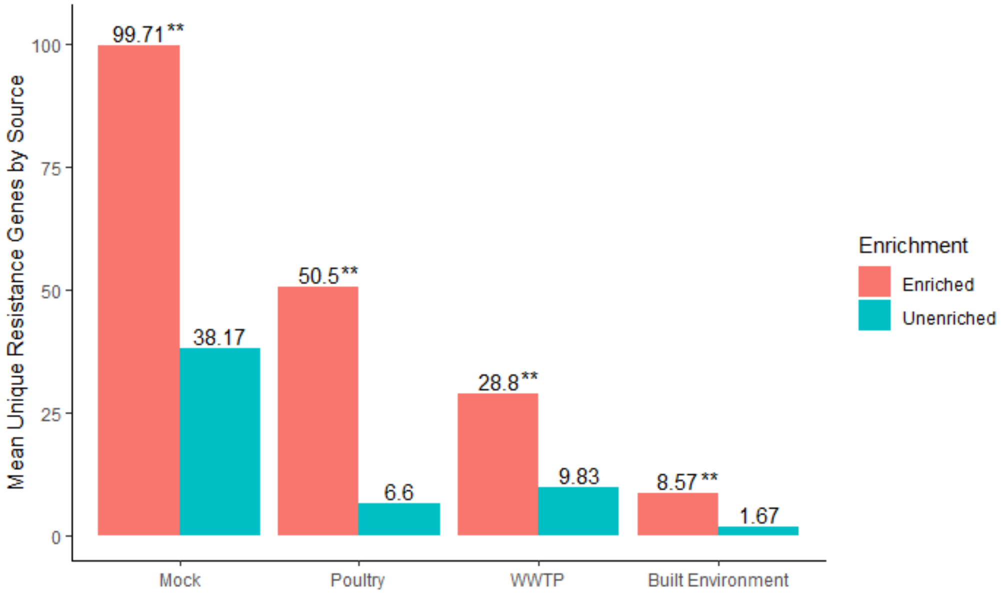


Supplemental Figure 2. Mean number of resistance genes broken down by sample type. Enriched libraries are red, unenriched libraries are teal. Stars represent a significant difference in the mean number of resistance genes between enriched and unenriched libraries measured by a Wilcoxon rank sum test (p-value ≤ 0.05).

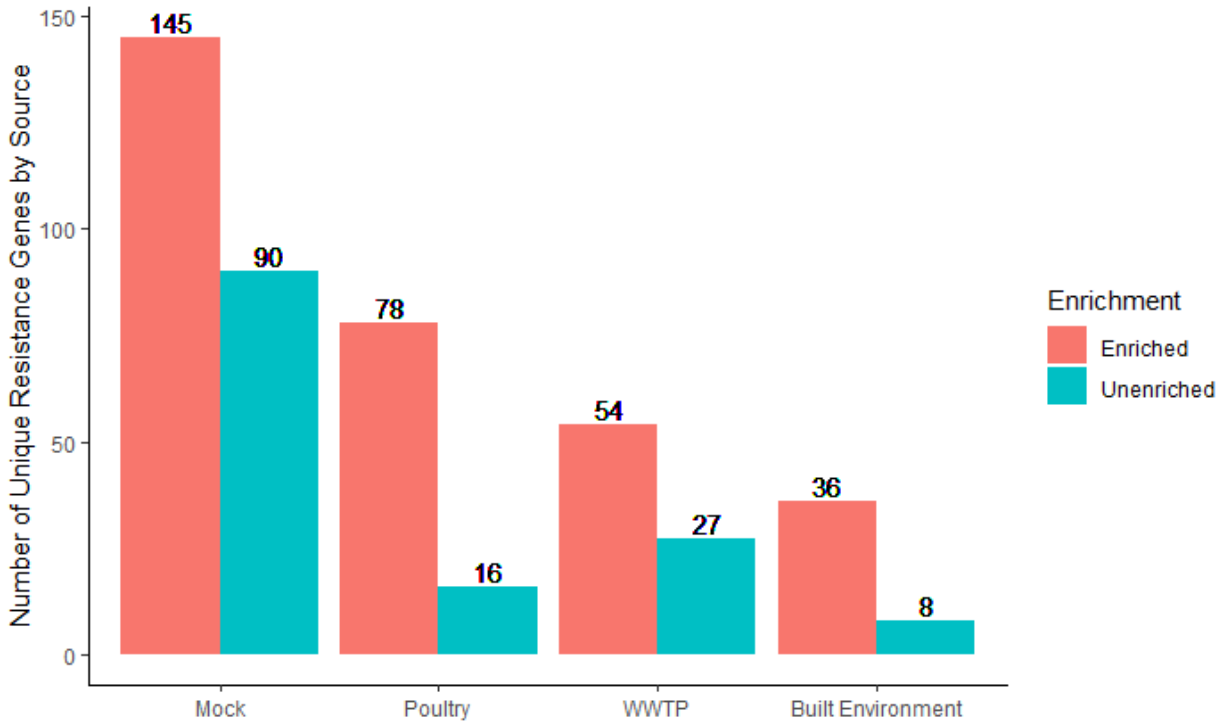


Supplemental Figure 3. Barplot of normalized number of unique resistance genes broken down by source (x-axis) and library type (i.e., enriched or unenriched), y-axis is number of unique resistance genes by source.

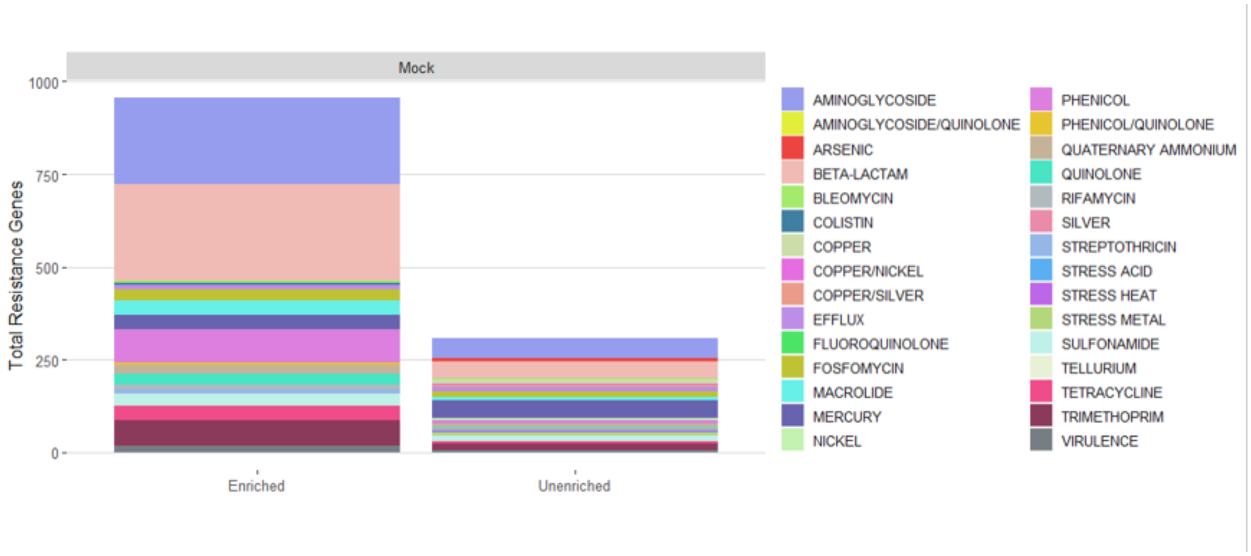


Supplemental Figure 4. Non-equalized de-novo analysis of the resistance classes in the mock community broken down by resistance class. Total resistance genes are on the y-axis and enriched and unenriched library types are on the x-axis.

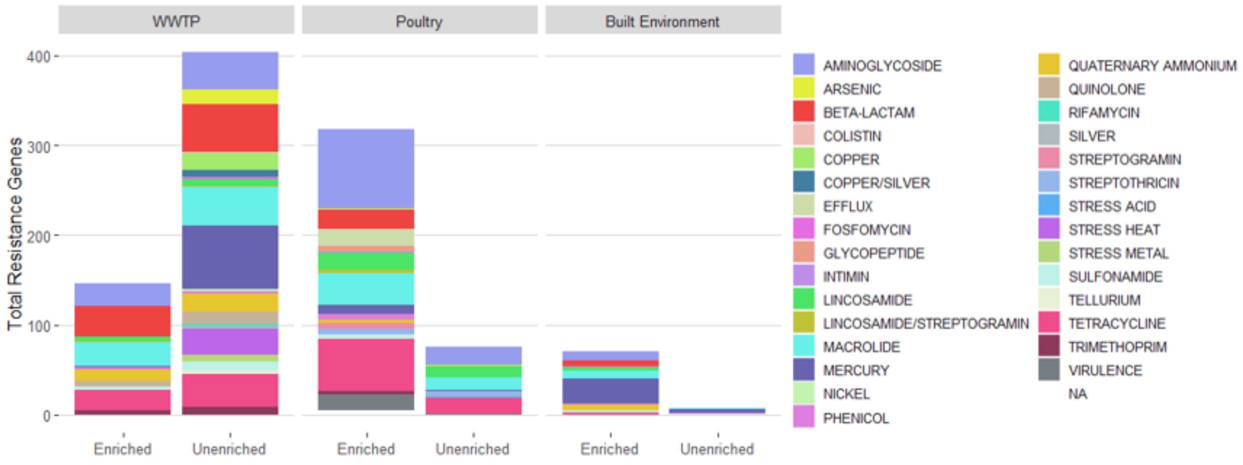


Supplemental Figure 5. Non-equalized de-novo analysis of the resistance classes in the environmental samples broken down by resistance class. Total resistance genes are on the y-axis and enriched and unenriched library types are on the x-axis.

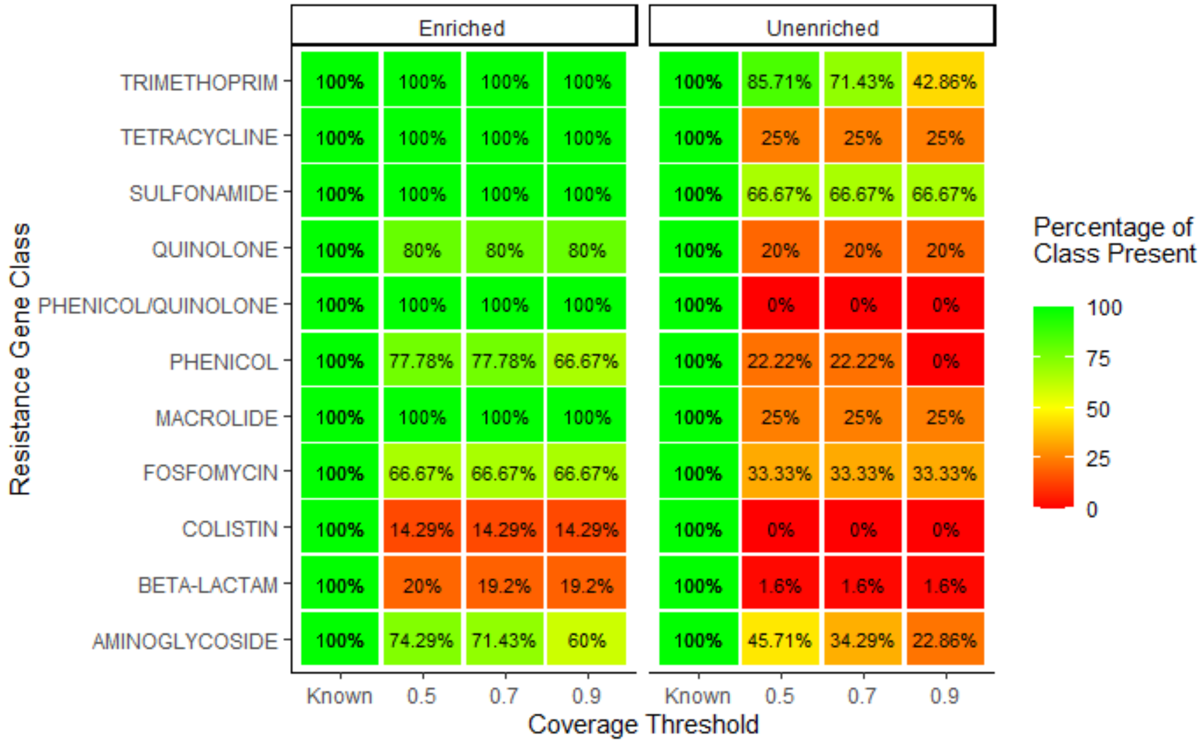


Supplemental Figure 6. Resistance mock community data analyzed at three different thresholds (i.e., 0.5, 0.7 and 0.9) in AMRfinder.

Supplemental Table 1. Antimicrobial resistance genes included in the mock community.

| Allele | Variant | Antibiotic Class | Gene Bank Accession | Product Annotation |
|---|---|---|---|---|
| **aac(2')-Ia** | aac(2')-Ia_1 | Aminoglycoside resistance | L06156 | aminoglycoside 2'-N-acetyltransferase |
| **aac(3)-Ia** | aac(3)-Ia_1 | Aminoglycoside resistance | X15852 | unnamed protein product; aacC1 product acetyltransferase (AA 1-177)" |
| **aac(3)-Ib** | aac(3)-Ib_1 | Aminoglycoside resistance | L06157 | aminoglycoside 3'-N-acetyltransferase |
| **aac(3)-Ic** | aac(3)-Ic_1 | Aminoglycoside resistance | AJ511268 | aminoglycoside acetyltransferase |
| **aac(3)-Id** | aac(3)-Id_1 | Aminoglycoside resistance | AB114632 | aminoglycoside acetyltransferase |
| **aac(3)-IIa** | aac(3)-IIa_1 | Aminoglycoside resistance | X51534 | unnamed protein product; AAC(3)-II (AA 1-286) |
| **aac(3)-IId** | aac(3)-IId_1 | Aminoglycoside resistance | EU022314 | aminoglycoside-(3)-N-acetyl-transferase |
| **aac(6')-33** | aac(6')-33_1 | Aminoglycoside resistance | GQ337064 | aminoglycoside acetyltransferase |
| **aac(6')-aph(2'')** | aac(6')-aph(2'')_1 | Aminoglycoside resistance | M13771 | AAC(6')-APH(2') bifunctional resistance protein |
| **aac(6')-Ib** | aac(6')-Ib_1 | Aminoglycoside resistance | M21682 | aminoglycoside 6'-N-acetyltransferase |
| **aac(6')Ib-cr** | aac(6')Ib-cr_1 | Fluoroquinolone and aminoglycoside resistance | DQ303918 | fluroquinolone acetylating aminoglycoside acetyltransferase |
| **aac(6')-Ic** | aac(6')-Ic_1 | Aminoglycoside resistance | M94066 | aminoglycoside acetyltransferase |
| **aac(6')-If** | aac(6')-If_1 | Aminoglycoside resistance | X55353 | AG-6'-acetyltransferase |
| **aac(6')-IIa** | aac(6')-IIa_1 | Aminoglycoside resistance | M29695 | 6'-N-acetyltransferase |
| **aac(6')-IIc** | aac(6')-IIc_1 | Aminoglycoside resistance | AF162771 | aminoglycoside 6'-N-acetyltransferase |
| **aadA1** | aadA1_4 | Aminoglycoside resistance | M95287 | aminoglycoside (3' ') adenylyltransferase |
| **aadA1** | aadA1_5 | Aminoglycoside resistance | JQ480156 | aminoglycoside-adenyltransferase |
| **aadA11** | aadA11_1 | Aminoglycoside resistance | AY144590 | aminoglycoside adenylyltransferase |
| **aadA2** | aadA2_2 | Aminoglycoside resistance | JQ364967 | aminoglycosides adenyltransferase |

| Allele | Variant | Antibiotic Class | Gene Bank Accession | Product Annotation |
|---|---|---|---|---|
| **aadA5** | aadA5_1 | Aminoglycosid e resistance | AF137361 | streptomycin and spectinomycin resistance aminoglycoside adenyltransferase |
| **aadA6** | aadA6_1 | Aminoglycoside resistance | AF140629 | aminoglycoside adenylyltransferase AADA6 |
| **aadA7** | aadA7_1 | Aminoglycoside resistance | AF224733 | aminoglycoside adenylyltransferase |
| **aadB** | aadB_1 | Aminoglycoside resistance | JN119852 | aminoglycoside-2"-adenylyltransferase" |
| **aadD** | aadD_1 | Aminoglycoside resistance | AF181950 | kanamycin resistance protein |
| **ACT-15** | ACT-15_1 | Beta-lactam resistance:AmpC - type | JX440356 | AmpC beta-lactamase ACT- 15 |
| **ACT-16** | ACT-16_1 | Beta-lactam resistance:AmpC - type | AB737978 | beta lactamase ACT-16 |
| **ACT-5** | ACT-5_1 | Beta-lactam resistance:AmpC - type | FJ237369 | beta-lactamase ACT-5 |
| **ACT-7** | ACT-7_1 | Beta-lactam resistance:AmpC-type | FJ237368 | beta-lactamase EBC-1464 |
| **ADC-25** | ADC-25_1 | Beta-lactam resistance | EF016355 | AmpC cephalosporinase |
| **aph(3')-Ia** | aph(3')-Ia_1 | Aminoglycoside resistance | V00359 | |
| **aph(3')-Ic** | aph(3')-Ic_1 | Aminoglycoside resistance | X62115 | neomycin phosphotransferase |
| **aph(3')-IIb** | aph(3')-IIb_1 | Aminoglycoside resistance | X90856 | aminoglycoside phosphotransferase (3')IIps |
| **aph(3')-III** | aph(3')-III_1 | Aminoglycoside resistance | M26832 | kanamycin resistance protein |
| **aph(3')-VIa** | aph(3')-VIa_1 | Aminoglycoside resistance | X07753 | |
| **aph(3')-XV** | aph(3')-XV_1 | Aminoglycoside resistance | Y18050 | aminoglycoside phosphotransferase |
| **aph(4)-Ia** | aph(4)-Ia_1 | Aminoglycoside resistance | V01499 | unnamed protein product; aph(4) |

| | | | | |
|---|---|---|---|---|
| **armA** | armA_1 | Aminoglycoside resistance | AY220558 | 16S rRNA methylase |
| **ARR-3** | ARR-3_4 | Rifampicin resistance | FM207631 | rifampicin ADP-ribosylatin transferase |

| Allele | Variant | Antibiotic Class | Gene Bank Accession | Product Annotation |
|---|---|---|---|---|
| **blaZ** | blaZ_34 | Beta-lactam resistance | AP003139 | beta-lactamase |
| **blaZ** | blaZ_35 | Beta-lactam resistance | AJ302698 | beta-lactamase |
| **blaZ** | blaZ_36 | Beta-lactam resistance | AJ400722 | beta-lactamase |
| **catA1** | catA1_1 | Phenicol resistance | V00622 | |
| **catA3** | catA3_1 | Phenicol resistance | X07848 | |
| **catB2** | catB2_1 | Phenicol resistance | AF047479 | chloramphenicol acetyltransferase |
| **catB3** | catB3_1 | Phenicol resistance | AJ009818 | Chloramphenicol acetyltransferase |
| **catB7** | catB7_1 | Phenicol resistance | AF036933 | chloramphenicol acetyltransferase |
| **catB8** | catB8_1 | Phenicol resistance | AF227506 | chloramphenicol acetyltransferase |
| **cmlA1** | cmlA1_1 | Phenicol resistance | M64556 | nonenzymatic chloramphenicol-resistance protein |
| **cmlA1** | cmlA1_2 | Phenicol resistance | AB212941 | chloramphenicol resistance |
| **cmx** | cmx_1 | Phenicol resistance | U85507 | chloramphenicol resistance protein |
| **CMY-2** | CMY-2_1 | Beta-lactam resistance | X91840 | extended spectrum beta-lactamase |
| **CMY-4** | CMY-4_1 | Beta-lactam resistance | AF420597 | cephamycinase |
| **CMY-42** | CMY-42_1 | Beta-lactam resistance | HM146927 | AmpC beta-lactamase |
| **CMY-6** | CMY-6_1 | Beta-lactam resistance | AJ011293 | beta-lactamase |
| **CMY-76** | CMY-76_1 | Beta-lactam resistance | JQ733573 | AmpC beta-lactamase CMY-76 |
| **CMY-79** | CMY-79_1 | Beta-lactam resistance | JQ733576 | AmpC beta-lactamase CMY-79 |
| **CMY-80** | CMY-80_1 | Beta-lactam resistance | JQ733577 | AmpC beta-lactamase CMY-80 |

| | | | | |
|---|---|---|---|---|
| **CMY-84** | CMY-84_1 | Beta-lactam resistance | JQ733579 | AmpC beta-lactamase CMY-84 |
| **CMY-94** | CMY-94_1 | Beta-lactam resistance | JX514368 | CMY-like beta-lactamase |

| Allele | Variant | Antibiotic Class | Gene Bank Accession | Product Annotation |
|---|---|---|---|---|
| **CTX-M-124** | CTX-M-124_1 | Beta-lactam resistance | JQ429324 | extended-spectrum beta-lactamase CTX-M-124 |
| **CTX-M-14** | CTX-M-14_1 | Beta-lactam resistance | AF252622 | beta-lactamase CTX-M-14 |
| **CTX-M-15** | CTX-M-15_23 | Beta-lactam resistance | DQ302097 | extended-spectrum beta-lactamase CTX-M-15 |
| **CTX-M-2** | CTX-M-2_1 | Beta-lactam resistance | EU622041 | resistance to beta-lactams including extended-spectrum cephalosporins |
| **CTX-M-55** | CTX-M-55_2 | Beta-lactam resistance | GQ456159 | extended-spectrum beta-lactamase CTX-M-55 |
| **dfrA1** | dfrA1_1 | Trimethoprim resistance | X00926 | dihydrofolate reductase (DHFR) |
| **dfrA1** | dfrA1_30 | Trimethoprim resistance | JQ690541 | dihydrofolate reductase type I |
| **dfrA12** | dfrA12_1 | Trimethoprim resistance | AB571791 | dihydrofolate reductase |
| **dfrA14** | dfrA14_1 | Trimethoprim resistance | DQ388123 | dihydrofolate reductase |
| **dfrA15** | dfrA15_1 | Trimethoprim resistance | HM449019 | dihydrofolate reductase |
| **dfrA17** | dfrA17_1 | Trimethoprim resistance | FJ460238 | dihydrofolate reductase; trimethoprim resistance |
| **dfrA18** | dfrA18_1 | Trimethoprim resistance | AJ310778 | dihydrofolate reductase |
| **dfrA29** | dfrA29_1 | Trimethoprim resistance | AM237806 | dihydrofolate reductase type VII |
| **dfrA30** | dfrA30_1 | Trimethoprim resistance | AM997279 | dihydrofolate reductase |
| **dfrA3b** | dfrA3b_1 | Trimethoprim resistance | AY878717 | dihydrofolate reductase type IIIb |
| **dfrA5** | dfrA5_1 | Trimethoprim resistance | X12868 | |
| **dfrA8** | dfrA8_1 | Trimethoprim resistance | U10186 | dihydrofolate reductase type VIII |
| **dfrB1** | dfrB1_2 | Trimethoprim resistance | U36276 | dihydrofolate reductase |
| **dfrB5** | dfrB5_1 | Trimethoprim resistance | AY943084 | dihydrofolate reductase |

| | | | | |
|---|---|---|---|---|
| **dfrG** | dfrG_1 | Trimethoprim resistance | AB205645 | dihydrofolate reductase |
| **DHA-1** | DHA-1_1 | Beta-lactam resistance:AmpC - type | Y16410 | beta-lactamase class C |

| Allele | Variant | Antibiotic Class | Gene Bank Accession | Product Annotation |
|---|---|---|---|---|
| **ere(A)** | ere(A)_2 | Macrolide resistance | AF099140 | erythromycin esterase |
| **erm(42)** | erm(42)_2 | Macrolide resistance | AB601890 | erythromycin resistance protein |
| **erm(A)** | erm(A)_1 | Macrolide resistance | X03216 | |
| **erm(B)** | erm(B)_1 | Macrolide resistance | JN899585 | rRNA methylase |
| **fosA** | fosA_10 | Fosfomycin resistance | EU195449 | putative fosfomycin-resistance protein |
| **fosA** | fosA_13 | Fosfomycin resistance | EU487198 | fosfomycin-resistance protein |
| **fosA** | fosA_14 | Fosfomycin resistance | AB522970 | fosfomycin resistance protein FosA3 |
| **fosA** | fosA_3 | Fosfomycin resistance | NZ_ACWO01000079 | |
| **fosA** | fosA_7 | Fosfomycin resistance | NZ_AFBO01000747 | |
| **GES-1** | GES-1_1 | Beta-lactam resistance | HQ170511 | extended-spectrum beta-lactamase GES-7 |
| **IMP-1** | IMP-1_1 | Beta-lactam resistance | DQ522237 | metallo-beta-lactamase IMP-1-like |
| **IMP-14** | IMP-14_1 | Beta-lactam resistance | AY553332 | metallo-beta-lactamase; carbapenemase IMP-14 |
| **IMP-4** | IMP-4_1 | Beta-lactam resistance | AF244145 | carbapenem-hydrolysing beta-lactamase |
| **KPC-2** | KPC-2_1 | Beta-lactam resistance | AY034847 | carbapenemase |
| **KPC-3** | KPC-3_1 | Beta-lactam resistance | HM769262 | beta-lactamase KPC-3 |
| **KPC-4** | KPC-4_1 | Beta-lactam resistance | FJ473382 | beta-lactamase KPC-4 |
| **KPC-5** | KPC-5_1 | Beta-lactam resistance | EU400222 | beta-lactamase KPC-5 |
| **KPC-6** | KPC-6_1 | Beta-lactam resistance | EU555534 | beta-lactamase KPC-6 |

| | | | | |
|---|---|---|---|---|
| **LEN16** | LEN16_1 | Beta-lactam resistance | AY743416 | LEN-type penicillinase |
| **MAL-1** | MAL-1_1 | Beta-lactam resistance | AJ277209 | beta-lactamase MAL-1 |
| **MAL-1** | MAL-1_2 | Beta-lactam resistance | AJ609506 | class A beta-lactamase |
| **mcr-1** | mcr-1_1 | Colistin resistance | KP347127 | phosphoethanolamine transferase |

| Allele | Variant | Antibiotic Class | Gene Bank Accession | Product Annotation |
|---|---|---|---|---|
| **mecA** | mecA_14 | Beta-lactam resistance | AB505630 | penicillin-binding protein 2' |
| **mecA** | mecA_15 | Beta-lactam resistance | AB505628 | penicillin-binding protein 2' |
| **mecA** | mecA_6 | Beta-lactam resistance | FR821779 | beta-lactam-inducible penicillin-binding protein |
| **mph(A)** | mph(A)_1 | Macrolide resistance | D16251 | macrolide 2'-phosphotransferase I |
| **mph(C)** | mph(C)_2 | Macrolide resistance | AF167161 | |
| **mph(E)** | mph(E)_3 | Macrolide resistance | EU294228 | macrolide 2'-phosphotransferase |
| **msr(A)** | msr(A)_3 | Macrolide, Lincosamide and Streptogramin B resistance | M81802 | 21kDa product of the C-terminal of MsrB |
| **msr(E)** | msr(E)_4 | Macrolide, Lincosamide and Streptogramin B resistance | EU294228 | |
| **NDM-1** | NDM-1_1 | Beta-lactam resistance | FN396876 | metallo-beta-lactamase |
| **NDM-5** | NDM-5_1 | Beta-lactam resistance | JN104597 | NDM-5 metallo-beta-lactamase |
| **NDM-6** | NDM-6_1 | Beta-lactam resistance | JN967644 | NDM carbapenemase |
| **NDM-7** | NDM-7_1 | Beta-lactam resistance | JX262694 | NDM-7 metallo-beta-lactamase |
| **NMC-A** | NMC-A_1 | Beta-lactam resistance | AJ536087 | class A nonmetallocarbapenamase |
| **OKP-B-2** | OKP-B-2_1 | Beta-lactam resistance | AM051151 | OKP-B beta-lactamase |

| | | | | |
|---|---|---|---|---|
| **oqxA** | oqxA_1 | Quinolone resistance | EU370913 | component of RND-type multidrug efflux pump that confers resistance to olaquindox |
| **oqxB** | oqxB_1 | Quinolone resistance | EU370913 | component of RND-type multidrug efflux pump that confers resistance to olaquindox |
| **OXA-1** | OXA-1_1 | Beta-lactam resistance | J02967 | OXA-1 beta-lactamase |
| **OXA-10** | OXA-10_2 | Beta-lactam resistance | EU886981 | Beta-lactamase |

| Allele | Variant | Antibiotic Class | Gene Bank Accession | Product Annotation |
|---|---|---|---|---|
| **OXA-100** | OXA-100_1 | Beta-lactam resistance | AM231720 | blaOXA-100 class D beta-lactamase |
| **OXA-181** | OXA-181_1 | Beta-lactam resistance | HM992946 | carbapenemase OXA-181 |
| **OXA-2** | OXA-2_1 | Beta-lactam resistance | DQ310703 | oxacillinase type 2 |
| **OXA-203** | OXA-203_1 | Beta-lactam resistance | HQ998857 | beta-lactamase OXA-203 |
| **OXA-223** | OXA-223_1 | Beta-lactam resistance | JN248564 | beta-lactamase OXA-223 |
| **OXA-23** | OXA-23_1 | Beta-lactam resistance | HQ700358 | beta-lactamase OXA-23 |
| **OXA-232** | OXA-232_1 | Beta-lactam resistance | JX423831 | class D beta-lactamase |
| **OXA-237** | OXA-237_1 | Beta-lactam resistance | JQ820241 | beta-lactamase OXA-237 |
| **OXA-24** | OXA-24_1 | Beta-lactam resistance | AJ239129 | Beta-lactamase |
| **OXA-4** | OXA-4_1 | Beta-lactam resistance | AY162283 | beta-lactamase OXA-4 |
| **OXA-48** | OXA-48_2 | Beta-lactam resistance | AY236073 | class D carbapenem-hydrolyzing beta-lactamase |
| **OXA-50** | OXA-50_1 | Beta-lactam resistance | AY306135 | oxacillinase |
| **OXA-50** | OXA-50_2 | Beta-lactam resistance | AY306133 | oxacillinase |
| **OXA-50** | OXA-50_3 | Beta-lactam resistance | AY306130 | oxacillinase |
| **OXA-50** | OXA-50_3 | Beta-lactam resistance | AY306132 | oxacillinase |

| | | | | |
|---|---|---|---|---|
| **OXA-56** | OXA-56_1 | Beta-lactam resistance | AY660529 | restricted spectrum class D beta-lactamase |
| **OXA-58** | OXA-58_1 | Beta-lactam resistance | AY665723 | carbapenem-hydrolyzing oxacillinase |
| **OXA-64** | OXA-64_1 | Beta-lactam resistance | AY750907 | beta-lactamase OXA-64 |
| **OXA-65** | OXA-65_1 | Beta-lactam resistance | AY750908 | beta-lactamase OXA-65 |
| **OXA-66** | OXA-66_4 | Beta-lactam resistance | FJ360530 | OXA carbapenemase; class D oxacillinase |
| **OXA-69** | OXA-69_1 | Beta-lactam resistance | HM564339 | carbapenem-hydrolyzing oxacillinase OXA-69 |

| Allele | Variant | Antibiotic Class | Gene Bank Accession | Product Annotation |
|---|---|---|---|---|
| **OXA-71** | OXA-71_1 | Beta-lactam resistance | AY859528 | carbapenem-hydrolyzing oxacillinase OXA-71 |
| **OXA-72** | OXA-72_1 | Beta-lactam resistance | GU199039 | class D beta-lactamase OXA-72 |
| **OXA-82** | OXA-82_1 | Beta-lactam resistance | GQ352402 | beta-lactamase OXA-82 protein |
| **OXA-9** | OXA-9_2 | Beta-lactam resistance | JF703130 | oxacillinase-carbenicillinase |
| **OXA-94** | OXA-94_1 | Beta-lactam resistance | DQ519088 | beta-lactamase OXA-94 |
| **OXY-1-4** | OXY-1-4_4 | Beta-lactam resistance | AY077483 | K1 beta-lactamase |
| **OXY-1-7** | OXY-1-7_7 | Beta-lactam resistance | M27459 | beta-lactamase |
| **OXY-2-8** | OXY-2-8_8 | Beta-lactam resistance | AY055205 | K1 beta-lactamase OXY-2 |
| **PAO** | PAO_1 | Beta-lactam resistance | AY083595 | beta-lactamase precursor |
| **PAO** | PAO_2 | Beta-lactam resistance | FJ666065 | class C beta-lactamase PDC-1 |
| **PAO** | PAO_3 | Beta-lactam resistance | FJ666073 | class C beta-lactamase PDC-10 |
| **PAO** | PAO_4 | Beta-lactam resistance | AY083592 | beta-lactamase precursor |
| **PER-7** | PER-7_1 | Beta-lactam resistance | HQ713678 | extended spectrum beta-lactamase |
| **QnrB13** | QnrB13_1 | Quinolone resistance | EU273755 | quinolone resistance protein; pentapeptide repeat protein |

| | | | | |
|---|---|---|---|---|
| **QnrB2** | QnrB2_1 | Quinolone resistance | HM125698 | fluoroquinolone resistance protein |
| **QnrB34** | QnrB34_1 | Quinolone resistance | JN173056 | quinolone resistance protein QnrB34 |
| **QnrB38** | QnrB38_1 | Quinolone resistance | JN173060 | QnrB38 |
| **QnrB4** | QnrB4_1 | Quinolone resistance | DQ303921 | contains pentapeptide repeat |
| **QnrB5** | QnrB5_1 | Quinolone resistance | DQ303919 | contains pentapeptide repeat |
| **QnrB6** | QnrB6_3 | Quinolone resistance | EF523819 | QnrB6 |
| **QnrB7** | QnrB7_1 | Quinolone resistance | EU043311 | QnrB7 |

| Allele | Variant | Antibiotic Class | Gene Bank Accession | Product Annotation |
|---|---|---|---|---|
| **QnrS1** | QnrS1_1 | Quinolone resistance | AB187515 | Qnr |
| **rmtB** | rmtB_1 | Aminoglycoside resistance | AB103506 | 16S rRNA methylase |
| **rmtC** | rmtC_1 | Aminoglycoside resistance | AB194779 | 16S rRNA methylase |
| **rmtf** | rmtf_1 | Aminoglycoside resistance | JQ808129 | aminoglycoside 16s rRNA methyltransferase |
| **rmtG** | rmtG_1 | Aminoglycoside resistance | KJ004567 | 16S rRNA methyltransferase |
| **SFO-1** | SFO-1_1 | Beta-lactam resistance | AB003148 | class A beta-lactamase SFO-1 |
| **SHV-1** | SHV-1_11 | Beta-lactam resistance | EF035565 | extended spectrum beta lactamase |
| **SHV-1** | SHV-1_13 | Beta-lactam resistance | AF462396 | extended-spectrum beta-lactamase SHV-1 |
| **SHV-1** | SHV-1_18 | Beta-lactam resistance | FJ668818 | beta-lactamase SHV-1 |
| **SHV-1** | SHV-1_9 | Beta-lactam resistance | HM751102 | SHV-1 beta-lactamase |
| **SHV-11** | SHV-11_10 | Beta-lactam resistance | EF035557 | extended spectrum beta lactamase |
| **SHV-11** | SHV-11_15 | Beta-lactam resistance | DQ219473 | extended-spectrum beta-lactamase SHV-11 |
| **SHV-11** | SHV-11_1 | Beta-lactam resistance | AY528717 | beta-lactamase |
| **SHV-11** | SHV-11_2 | Beta-lactam resistance | HM751098 | SHV-11 beta-lactamase |

| | | | | |
|---|---|---|---|---|
| **SHV-11** | SHV-11_5 | Beta-lactam resistance | GQ407117 | beta-lactamase blaSHV-11 |
| **SHV-11** | SHV-11_8 | Beta-lactam resistance | GQ387358 | beta-lactamase SHV-11 |
| **SHV-11** | SHV-11_9 | Beta-lactam resistance | FJ483937 | beta-lactamase SHV-11 |
| **SHV-12** | SHV-12_1 | Beta-lactam resistance | AF462395 | extended-spectrum beta-lactamase SHV-12 |
| **SHV-26** | SHV-26_1 | Beta-lactam resistance | AF227204 | beta-lactamase |
| **SHV-30** | SHV-30_1 | Beta-lactam resistance | AY661885 | beta-lactamase SHV-30 |
| **SHV-5** | SHV-5_6 | Beta-lactam resistance | EF653399 | beta-lactamase SHV-5 |
| **SHV-83** | SHV-83_1 | Beta-lactam resistance | AM176558 | beta-lactamase |

| Allele | Variant | Antibiotic Class | Gene Bank Accession | Product Annotation |
|---|---|---|---|---|
| **SME-3** | SME-3_1 | Beta-lactam resistance | AY584237 | carbapenem-hydrolyzing beta-lactamase SME-3 |
| **spc** | spc_1 | Aminoglycoside resistance | X02588 | |
| **SPM-1** | SPM-1_1 | Beta-lactam resistance | AY341249 | carbapenemase |
| **strA** | strA_1 | Aminoglycoside resistance | M96392 | |
| **strA** | strA_4 | Aminoglycoside resistance | AF321551 | aminoglycoside 3"-phosphotransferase |
| **strB** | strB_1 | Aminoglycoside resistance | M96392 | streptomycin phosphotransferase |
| **sul1** | sul1_11 | Sulphonamide resistance | DQ914960 | dihydropteroate synthase |
| **sul1** | sul1_2 | Sulphonamide resistance | CP002151 | dihydropteroate synthase protein Sul1 |
| **sul2** | sul2_2 | Sulphonamide resistance | GQ421466 | dihydropteroate synthase |
| **sul2** | sul2_3 | Sulphonamide resistance | HQ840942 | sulfonamide-insensitive dihydropteroate synthetase |
| **sul3** | sul3_2 | Sulphonamide resistance | AJ459418 | dihydropteroate synthase |
| **TEM-1A** | TEM-1A_4 | Beta-lactam resistance | HM749966 | TEM-1 beta-lactamase |
| **TEM-1B** | TEM-1B_1 | Beta-lactam resistance | JF910132 | beta-lactamase |

| | | | | |
|---|---|---|---|---|
| **TEM-1D** | TEM-1D_83 | Beta-lactam resistance | AF188200 | beta-lactamase variant TEM-1D |
| **TEM-52B** | TEM-52B_1 | Beta-lactam resistance | AF027199 | extended spectrum beta-lactamase |
| **tet(A)** | tet(A)_3 | Tetracycline resistance | AY196695 | tetracycline resistance efflux pump |
| **tet(A)** | tet(A)_4 | Tetracycline resistance | AJ517790 | tetracycline resistance protein |
| **tet(B)** | tet(B)_3 | Tetracycline resistance | AP000342 | tetracycline resistance protein |
| **tet(B)** | tet(B)_4 | Tetracycline resistance | AF326777 | tetracycline resistance protein TetA(B) |
| **tet(C)** | tet(C)_5 | Tetracycline resistance | NC_003213 | TetA(C) protein |
| **tet(D)** | tet(D)_1 | Tetracycline resistance | AF467077 | TetA(D); class D; efflux pump |

| Allele | Variant | Antibiotic Class | Gene Bank Accession | Product Annotation |
|---|---|---|---|---|
| **tet(G)** | tet(G)_4 | Tetracycline resistance | AF133140 | tetracycline resistance protein |
| **tet(J)** | tet(J)_1 | Tetracycline resistance | ACLE01000065 | transporter, major facilitator family protein |
| **tet(K)** | tet(K)_4 | Tetracycline resistance | U38428 | tetracycline resistance protein |
| **tet(M)** | tet(M)_7 | Tetracycline resistance | FN433596 | tetracycline resistance protein |
| **VEB-1** | VEB-1_1 | Beta-lactam resistance | HM370393 | extended-spectrum beta-lactamase |
| **VIM-1** | VIM-1_1 | Beta-lactam resistance | Y18050 | beta-lactamase VIM-1 |
| **VIM-11** | VIM-11_1 | Beta-lactam resistance | AY605049 | VIM-11 |
| **VIM-2** | VIM-2_1 | Beta-lactam resistance | AF302086 | metallo-beta-lactamase |
| **VIM-27** | VIM-27_1 | Beta-lactam resistance | HQ858608 | metallo-beta-lactamase VIM-27 |
| **VIM-4** | VIM-4_1 | Beta-lactam resistance | EU581706 | metallo-beta-lactamase |

Supplemental Table 2. Outputs from CARDv1.0 not mapping to baits designed in this study (i.e., AMR-Cap baits) including accession and gene.

| Accession | Gene |
|---|---|
| ARO:3002813 | lmrB |
| ARO:3000834 | phoP |

| | |
|---|---|
| ARO:3003582 | PmrA |
| ARO:3000198 | FosX |
| ARO:3000822 | pmrA |
| ARO:3003552 | fusB |
| ARO:3000309 | emrD |
| ARO:3000621 | PC1 |
| ARO:3000621 | PC1 |
| ARO:3000863 | mprF |
| ARO:3003043 | PBP2x |
| ARO:3003041 | PBP1a |
| ARO:3003098 | cphA1 |
| ARO:3000818 | nalC |
| ARO:3003044 | PBP1b |
| ARO:3001327 | mdtK |
| ARO:3000835 | phoQ |
| ARO:3003042 | PBP2b |

Supplementary Table 3. Outputs from CARDv3.3.1 not mapping to baits designed in this study (i.e., AMR-Cap baits) including accession and gene.

| Accession | Gene |
|---|---|
| ARO:3005084 | dfrA31 |
| ARO:3004638 | AAC(6')-Iag |
| ARO:3004768 | CIA-1 |
| ARO:3004628 | AAC(2')-IIa |
| ARO:3004769 | CIA-2 |
| ARO:3004770 | CIA-3 |
| ARO:3004771 | CIA-4 |
| ARO:3004106 | TMB-2 |
| ARO:3004588 | Klebsiella |
| ARO:3004289 | Vibrio |
| ARO:3004364 | almG |
| ARO:3004077 | PmpM |
| ARO:3005067 | ParS |
| ARO:3005068 | ParR |
| ARO:3004107 | Pseudomonas |
| ARO:3004072 | OpmB |
| ARO:3004075 | MuxC |
| ARO:3004074 | MuxB |
| ARO:3004073 | MuxA |
| ARO:3005063 | cprR |

| | |
|---|---|
| ARO:3005064 | cprS |
| ARO:3004056 | ArmR |
| ARO:3000025 | patB |
| ARO:3000024 | patA |
| ARO:3005045 | SatA |
| ARO:3005100 | FosB2 |
| ARO:3004466 | ICR-Mc |
| ARO:3004784 | FAR-1 |
| ARO:3005069 | rsmA |
| ARO:3004680 | APH(3')-VIIIa |
| ARO:3004751 | BES-1 |
| ARO:3004659 | Mef(En2) |
| ARO:3004740 | AST-1 |
| ARO:3004448 | HERA-1 |
| ARO:3004766 | CGA-1 |
| ARO:3005008 | Txr |
| ARO:3004817 | HERA-2 |
| ARO:3004818 | HERA-3 |
| ARO:3004669 | emtA |
| ARO:3004780 | DES-1 |
| ARO:3000168 | tet(D) |
| ARO:3004773 | CKO-1 |
| ARO:3004661 | Staphylococcus |
| ARO:3002875 | dfrE |
| ARO:3004775 | CME-1 |
| ARO:3004359 | ACI-1 |
| ARO:3004122 | Klebsiella |
| ARO:3004788 | FONA-1 |
| ARO:3004789 | FONA-2 |
| ARO:3004790 | FONA-3 |
| ARO:3004791 | FONA-4 |
| ARO:3004792 | FONA-5 |
| ARO:3004793 | FONA-6 |
| ARO:3005166 | Tet(X1) |
| ARO:3004819 | HERA-4 |
| ARO:3004820 | HERA-5 |
| ARO:3004821 | HERA-6 |
| ARO:3004822 | HERA-8 |
| ARO:3004747 | BAT-1 |
| ARO:3003071 | mphF |

| | |
|---|---|
| ARO:3003854 | ADC-8 |
| ARO:3004146 | cfrC |
| ARO:3004605 | ermZ |
| ARO:3000501 | Nocardia |
| ARO:3004580 | Klebsiella |
| ARO:3004583 | Klebsiella |
| ARO:3004612 | Escherichia |
| ARO:3005030 | FRI-7 |
| ARO:3005032 | FRI-9 |
| ARO:3005031 | FRI-8 |
| ARO:3001741 | OXA-286 |
| ARO:3004091 | ANT(3")-IIc |
| ARO:3001734 | OXA-279 |
| ARO:3001749 | OXA-294 |
| ARO:3004090 | ANT(3")-IIb |
| ARO:3004087 | APH(3')-IX |
| ARO:3001763 | OXA-308 |
| ARO:3001722 | OXA-266 |
| ARO:3001754 | OXA-299 |
| ARO:3001746 | OXA-291 |
| ARO:3001743 | OXA-288 |
| ARO:3001747 | OXA-292 |
| ARO:3001752 | OXA-297 |
| ARO:3001757 | OXA-302 |
| ARO:3001750 | OXA-295 |
| ARO:3001753 | OXA-298 |
| ARO:3001761 | OXA-306 |
| ARO:3001758 | OXA-303 |
| ARO:3004479 | OXA-664 |
| ARO:3001730 | OXA-274 |
| ARO:3004086 | APH(3')-VIII |
| ARO:3004478 | OXA-665 |
| ARO:3004568 | dfrA18 |
| ARO:3004782 | ERP-1 |
| ARO:3004642 | dfrA3b |
| ARO:3004123 | Burkholderia |
| ARO:3005111 | blaS1 |
| ARO:3004292 | Laribacter |
| ARO:3005088 | msrH |
| ARO:3003733 | fusC |

| | |
|---|---|
| ARO:3002983 | amrB |
| ARO:3002982 | amrA |
| ARO:3004572 | Staphylococcus |
| ARO:3004291 | Rhodobacter |
| ARO:3004573 | Acinetobacter |
| ARO:3004645 | dfrI |
| ARO:3004759 | BPU-1 |
| ARO:3004574 | Acinetobacter |
| ARO:3003725 | vanXI |
| ARO:3004801 | FTU-1 |
| ARO:3004539 | mphH |
| ARO:3005091 | RanA |
| ARO:3005090 | RanB |
| ARO:3005059 | LptD |
| ARO:3004795 | FPH-1 |
| ARO:3004541 | mphK |
| ARO:3003811 | adeC |
| ARO:3005162 | FosA7.5 |
| ARO:3004603 | tet(56) |
| ARO:3005051 | LpsB |
| ARO:3004099 | LpeA |
| ARO:3004100 | LpeB |
| ARO:3004667 | Staphylococcus |
| ARO:3005035 | Yrc-1 |
| ARO:3005052 | LpsA |
| ARO:3003741 | mphE |
| ARO:3004825 | LAP-1 |
| ARO:3004856 | SCO-1 |
| ARO:3004749 | BCL-1 |
| ARO:3002881 | lmrC |
| ARO:3004826 | LAP-2 |
| ARO:3004601 | LnuP |
| ARO:3004547 | dfrA6 |
| ARO:3004577 | Acinetobacter |
| ARO:3004691 | MCR-3.4 |
| ARO:3004551 | dfrA28 |
| ARO:3004611 | Escherichia |
| ARO:3002831 | vgaC |
| ARO:3005047 | eptB |
| ARO:3005044 | OmpA |

| | |
|---|---|
| ARO:3005053 | ArnT |
| ARO:3004105 | TMB-1 |
| ARO:3005046 | MecC-type |
| ARO:3004753 | BIC-1 |
| ARO:3004808 | GOB-16 |
| ARO:3004678 | rmtE |
| ARO:3005049 | CrcB |
| ARO:3003971 | erm(44) |
| ARO:3004666 | pexA |
| ARO:3004468 | CrpP |
| ARO:3004641 | aacA43 |
| ARO:3004033 | tetB(46) |
| ARO:3004032 | tetA(46) |
| ARO:3000510 | mupB |
| ARO:3004786 | FIM-1 |
| ARO:3004063 | EdeQ |
| ARO:3004542 | mphN |
| ARO:3001558 | OXA-372 |
| ARO:3004596 | erm(46) |
| ARO:3004550 | dfrA27 |
| ARO:3005006 | tet(57) |
| ARO:3004608 | EreD |
| ARO:3003746 | optrA |
| ARO:3004757 | BKC-1 |
| ARO:3004764 | CBP-1 |
| ARO:3001460 | OXA-155 |
| ARO:3001462 | OXA-157 |
| ARO:3000620 | adeL |
| ARO:3004613 | Tet(47) |
| ARO:3004581 | tet(48) |
| ARO:3004582 | tet(49) |
| ARO:3004584 | tet(50) |
| ARO:3004586 | tet(51) |
| ARO:3004587 | tet(52) |
| ARO:3004589 | tet(53) |
| ARO:3004590 | tet(54) |
| ARO:3004591 | tet(55) |
| ARO:3004797 | FRI-1 |
| ARO:3004679 | rmtE2 |
| ARO:3003620 | OXA-464 |

| | |
|---|---|
| ARO:3004441 | tet(59) |
| ARO:3004035 | tetA(60) |
| ARO:3004036 | tetB(60) |
| ARO:3004085 | lnuG |
| ARO:3003980 | tetA(48) |
| ARO:3003981 | tetB(48) |
| ARO:3003982 | LlmA |
| ARO:3003983 | CatU |
| ARO:3003984 | BahA |
| ARO:3003986 | TaeA |
| ARO:3003987 | VatI |
| ARO:3003988 | AAC(2')-IIb |
| ARO:3003992 | rphB |
| ARO:3003989 | AAC(6')-34 |
| ARO:3003994 | cpaA |
| ARO:3003990 | VgbC |
| ARO:3003991 | mphI |
| ARO:3004798 | FRI-2 |
| ARO:3004185 | mecD |
| ARO:3005150 | PAC-1 |
| ARO:3004737 | ARL-4 |
| ARO:3004735 | ARL-2 |
| ARO:3004736 | ARL-3 |
| ARO:3004738 | ARL-5 |
| ARO:3004734 | ARL-1 |
| ARO:3004739 | ARL-6 |
| ARO:3004799 | FRI-3 |
| ARO:3004544 | mphJ |
| ARO:3004357 | catV |
| ARO:3004332 | MCR-5 |
| ARO:3005164 | dfrA35 |
| ARO:3004139 | MCR-3 |
| ARO:3003060 | tsnR |
| ARO:3004113 | FosA7 |
| ARO:3005027 | FRI-4 |
| ARO:3004595 | erm45 |
| ARO:3004682 | aadA27 |
| ARO:3004094 | Erm(48) |
| ARO:3004054 | Pseudomonas |
| ARO:3001459 | OXA-154 |

| | |
|---|---|
| ARO:3004730 | tva(A) |
| ARO:3004699 | sta |
| ARO:3004683 | aadS |
| ARO:3004470 | poxtA |
| ARO:3004571 | PNGM-1 |
| ARO:3004505 | MCR-3.5 |
| ARO:3004662 | MCR-3.3 |
| ARO:3004325 | MCR-4 |
| ARO:3004503 | MCR-3.6 |
| ARO:3004510 | MCR-3.7 |
| ARO:3004509 | MCR-3.8 |
| ARO:3004508 | MCR-3.9 |
| ARO:3004695 | MCR-4.3 |
| ARO:3004504 | MCR-3.10 |
| ARO:3004517 | MCR-7.1 |
| ARO:3005033 | FLC-1 |
| ARO:3004698 | MCR-5.2 |
| ARO:3004559 | CAM-1 |
| ARO:3004500 | MCR-3.11 |
| ARO:3004693 | MCR-3.12 |
| ARO:3005021 | cfr(D) |
| ARO:3004516 | MCR-8 |
| ARO:3004445 | RSA-2 |
| ARO:3004444 | RSA-1 |
| ARO:3004694 | MCR-4.2 |
| ARO:3004697 | MCR-4.5 |
| ARO:3004696 | MCR-4.4 |
| ARO:3005028 | FRI-5 |
| ARO:3005096 | GPC-1 |
| ARO:3005121 | lsa(D) |
| ARO:3005018 | LMB-1 |
| ARO:3005029 | FRI-6 |
| ARO:3005060 | OXA-837 |
| ARO:3005061 | AAC(3)-IVb |
| ARO:3005062 | ANT(3")-Ib |
| ARO:3005024 | FosL1 |
| ARO:3005015 | VMB-1 |
| ARO:3005089 | mef(D) |
| ARO:3005087 | msrF |
| ARO:3005013 | IDC-2 |

| | |
|---|---|
| ARO:3005012 | IDC-1 |
| ARO:3005085 | AAC(3)-IIg |
| ARO:3004626 | Erm(49) |
| ARO:3004476 | vmlR |
| ARO:3004639 | Corynebacterium |
| ARO:3004480 | Bifidobacterium |
| ARO:3004684 | MCR-9 |
| ARO:3004643 | Erm(K) |
| ARO:3004102 | kamB |
| ARO:3001742 | OXA-287 |
| ARO:3001748 | OXA-293 |
| ARO:3004361 | sul4 |
| ARO:3004511 | MCR-3.2 |
| ARO:3004569 | ICR-Mo |
| ARO:3004290 | Escherichia |
| ARO:3004650 | tetU |
| ARO:3004635 | AAC(6')-Il |
| ARO:3004761 | BRO-1 |
| ARO:3004548 | dfrA9 |
| ARO:3000521 | mupA |
| ARO:3003836 | qacH |
| ARO:3004762 | BRO-2 |
| ARO:3004644 | dfrA6 |

Supplemental Table 4: Break down of predictions for CARD (signified by a *, assuming the same rate of gene acquisition) and the bait set designed in our study ability to capture novel resistance genes added to CARD.

| | CARDv1 | CARDv3 | CARDv5* | CARDv8* | CARDv10* |
|---|---|---|---|---|---|
| Total Resistance Genes | 2110 | 2786 | 3237 | 3913 | 5039 |
| New Resistance Genes | NA | 676 | 1127 | 1803 | 2253 |
| Years since CARDv1 | NA | 6 | 10 | 16 | 20 |
| Our baits: All resistance genes | 2091 | 2503 | 2765 | 3158 | 3420 |
| Our baits: New resistance genes | NA | 393 | 655 | 1048 | 1310 |
| Percent of Resistance Genes captured in CARD | 99.10% | 89.84% | 85.43% | 80.71% | 67.87% |
